# Receptor Tyrosine Kinase Profiling Identifies Chronic Constitutive Floodgate Oxidative Signaling in Glutathione-Independent Human Mammary Luminal Progenitor Cells

**DOI:** 10.1101/2025.07.15.665005

**Authors:** Syed Musheer Mohammed Aalam, Megan L. Ritting, Hari PS, Geng-Xian Shi, Kingsley Shih, Yifei Dong, Guruprasad Kalthur, Cao Miao, Yifan Yu, David JHF Knapp, Peter Eirew, Davide Pellacani, Chitra Priya Emperumal, Hequn Wang, Jeffrey R. Janus, Mark E. Sherman, Mathew P. Goetz, Sarah A. McLaughlin, Judy C. Boughey, Amy C. Degnim, Haishan Zeng, Akilesh Pandey, Derek C. Radisky, Anguraj Sadanandam, Nagarajan Kannan

**Author notes:** **Corresponding Author:** Nagarajan Kannan Ph.D., 200 First St, SW, Rochester, MN 55905.

## Abstract

The human mammary epithelium contains a subset of luminal progenitor (LP) cells that are distinct from basal cells in both lineage potential and redox biology. LPs are uniquely equipped to tolerate oxidative stress through glutathione-independent mechanisms and have been implicated as candidate cells of origin in basal-like breast cancers. In this study, we identify the receptor tyrosine kinase (RTK) cKIT (CD117), as a defining feature of LPs and a key mediator of their expansion. cKIT is developmentally restricted to the LP compartment via Polycomb-mediated epigenetic repression in basal and luminal-committed cells. It is expressed in scattered epithelial cells within both ductal and alveolar regions of resting human mammary glands.

Using RTK-engineered MCF10A models, we demonstrate that cKIT ligand/stem cell factor (SCF)-activated wildtype cKIT signaling is sufficient to drive proliferation in the absence of epidermal growth factor (EGF) and that cKIT is responsive not only to canonical ligands but also to hydrogen peroxide (H₂O₂). In primary human LPs, cKIT is rapidly phosphorylated upon exposure to SCF and H₂O₂, with concomitant AKT activation. These responses are enhanced when cKIT and EGFR signaling are co-engaged, suggesting a cooperative mitogenic program. In mammary gland, phosphorylation of the antioxidant enzyme PRDX1 is selectively detected in LPs, consistent with a floodgate model of redox signaling in which transient oxidative inactivation of peroxiredoxins (PRDXs) facilitates RTK signaling under elevated intracellular reactive oxygen species conditions.

Clinically, elevated cKIT expression is associated with shorter progression-free survival in certain basal-like breast cancer, supporting a link between LP-like redox signaling states and aggressive tumor behavior.

Together, these findings define a redox-integrated RTK signaling axis centered on cKIT that drives LP expansion and is associated with poor outcomes in a subset of basal breast cancers. This work establishes a mechanistic framework for targeting redox-responsive progenitor populations in both regenerative and oncologic context.

## INTRODUCTION

The human mammary epithelium consists of two principal cell lineages: basal myoepithelial and luminal epithelial cells. The basal compartment includes cells defined phenotypically as Lin⁻CD49f^high^EpCAM^low/⁻^ (basal cells, BCs), resident multipotent stem cells with mammary repopulating activity, myoepithelial-luminal bipotent progenitors and myoepithelial-restricted progenitors (1,2). The luminal compartment is composed of at least two functionally distinct populations: luminal progenitors (LPs; Lin⁻CD49f⁺EpCAM⁺), which display robust proliferative potential in culture, and luminal-committed (LC) cells (Lin⁻CD49f^low/⁻^EpCAM⁺), which are non-clonogenic under current assay conditions and are considered more differentiated (3). LPs are of particular interest because accumulating genetic and functional evidence implicates them as the likely cells of origin for basal-like breast cancers, particularly in individuals carrying BRCA1 mutations (2). Despite their importance, the upstream signaling pathways that regulate LP behavior, especially their proliferative response and resilience to oxidative stress, remain incompletely defined (1,2,4,5).

We previously showed that LPs, unlike BCs, retain proliferative capacity even when intracellular glutathione levels are depleted, suggesting that LPs rely on an alternative oxidative stress response mechanism (3). This redox flexibility is especially relevant in physiological states such as lactation, which imposes a sustained oxidative load on the mammary epithelium. Local accumulation of hydrogen peroxide (H₂O₂) during milk production can reach levels generally toxic to stem and progenitor cells in the basal epithelium, but are tolerated by progenitors in the luminal epithelium—a property that may be particularly relevant during lactation and could persist or be modulated during weaning and post-weaning involution, when oxidative remodeling continues (6). Notably, immunoglobulins, abundant in human milk, have been shown to catalyze the generation of reactive oxygen species, including H₂O₂, through antibody-mediated oxidation of water, potentially contributing to the luminal oxidative milieu during secretory activation(7). Recent single-cell transcriptomic analyses further support that LPs, not LCs, give rise to the secretory alveolar cells of the lactating gland (8). These observations underscore that LPs must not only proliferate under oxidative conditions but also do so in a tightly regulated manner.

Understanding how LPs integrate oxidative cues with lineage-specific mitogenic signaling is therefore central to unraveling the molecular logic of mammary epithelial homeostasis. Moreover, such insights may shed light on how aberrant redox signaling contributes to the early stages of breast cancer development, particularly in high-risk genetic contexts. This is characterized by deficient DNA repair mechanisms (e.g., *BRCA1* pathogenic variants) or precursors characterized by somatic alterations with similar loss of function. The “floodgate” hypothesis posits that overwhelming oxidative stresses may inactivate peroxiredoxins, especially PRDX1, which is abundant in epithelial cells thereby amplifying RTK signaling under oxidative stress (9). EGFR has been the primary model for this mechanism, but little is known about other RTKs that may be subject to redox modulation, particularly within lineage-restricted compartments of the human mammary gland.

In our study, we identified cKIT as a developmentally confined RTK, uniquely expressed in the CD49f⁺ luminal progenitor population. We demonstrated that cKIT is epigenetically restricted to LPs via Polycomb-mediated H3K27me3 repression in basal and LC populations. Functionally, cKIT integrates canonical (SCF) and redox (H₂O₂) signals to promote LP proliferation and cooperates with EGFR to amplify downstream mitogenic pathways. Importantly, LPs, but not basal cells (BCs) or luminal cells (LCs) exhibit constitutive phosphorylation of PRDX1, suggesting they operate under a tonic oxidative state that supports redox-sensitive kinase signaling.

## RESULTS

### cKIT Expression in the Mammary Epithelium is Restricted to Luminal Progenitors via Epigenetic Silencing

To define lineage-specific expression patterns of the cKIT receptor in the human mammary epithelium, we analyzed multiple transcriptomic datasets of fluorescence-activated cell sorting (FACS)-purified basal myoepithelial cells (BC; Lin⁻CD49f^high^EpCAM^low/^⁻), two luminal subpopulations namely progenitor-rich luminal cells (LP; Lin⁻CD49f⁺EpCAM⁺), and hormone-responsive luminal cells (LC; Lin⁻CD49f^low/^⁻EpCAM⁺) populations published by us (**GSE37223; EGAS00001001310**) (10,11) and others (**GSE16997, GSE17072**) (2). Both bulk epithelial subpopulation-specific RNA-sequencing and microarray datasets consistently demonstrated that *cKIT* transcripts were predominant RTKs in LP cells, while expression was markedly reduced or undetectable in LC and BC populations (**Figure 1A-C, Supplementary Figure S1A**). To validate our observations, we enzymatically dissociated human breast tissue to obtain single-cell suspensions and performed scRNA-seq, following protocols described previously (12). Consistent with our earlier findings, RTK profiling across epithelial subpopulations revealed distinct expression patterns. cKIT expression was predominantly confined to the LP clusters, with minimal to undetectable expression in BC or LC clusters (**Figure 1D-E**). In contrast, EGFR expression was enriched in both basal and LP populations (**Figure 1D, F**), underscoring the lineage specificity of cKIT within the mammary epithelial hierarchy.

**Figure 1.**
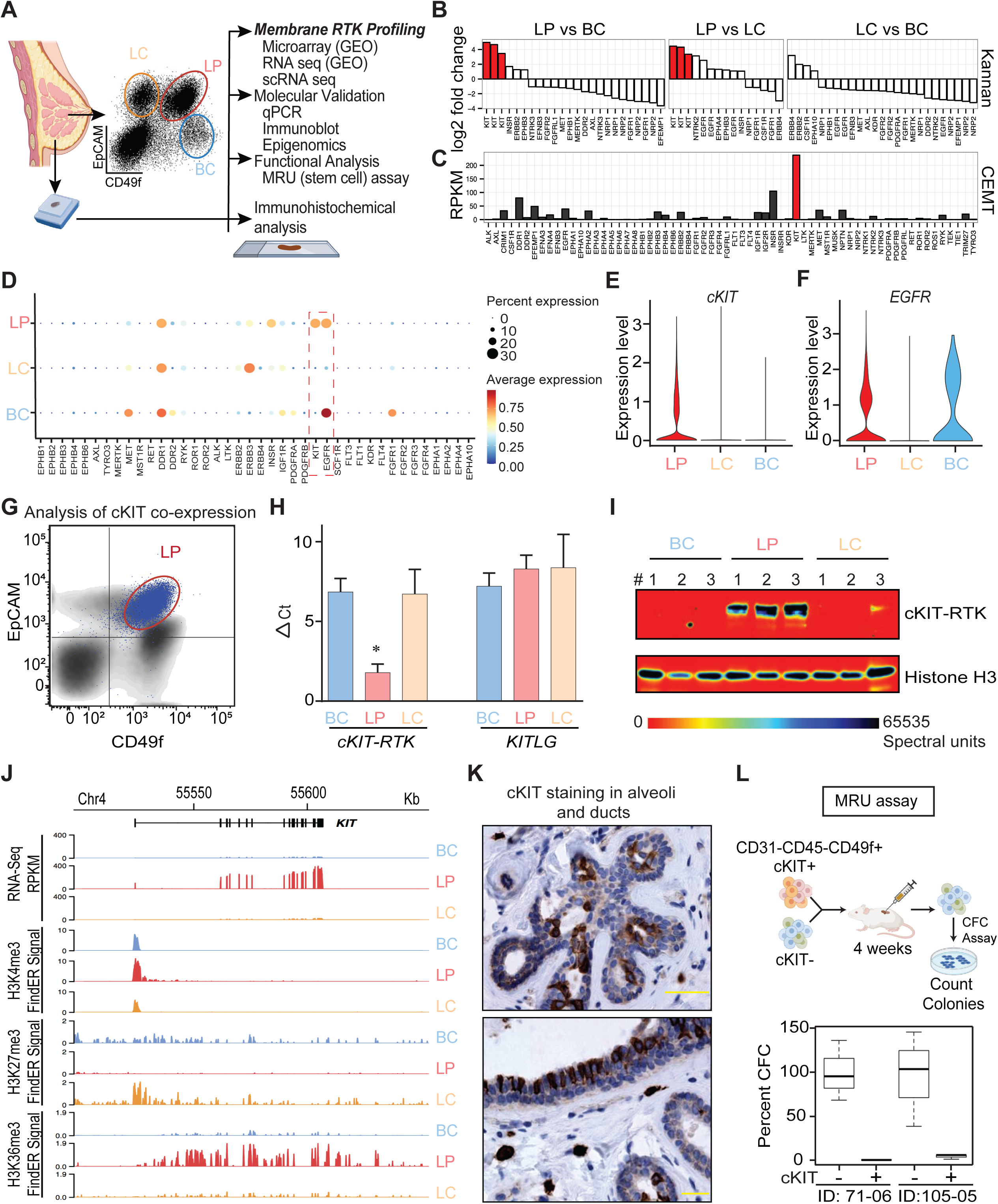
RTK Profiling Identifies cKIT as an Epigenetically Restricted RTK Enriched in Human Luminal Progenitor Cells. **(A)** Schematic overview of the integrated experimental workflow used to identify cKIT as a luminal progenitor (LP)-restricted receptor tyrosine kinase (RTK) in the human mammary epithelium. **(B–C)** Transcriptomic analysis of FACS-sorted epithelial subsets—basal cells (BCs: Lin⁻CD49f^high^EpCAM^low/⁻^), luminal progenitors (LPs: Lin⁻CD49f⁺EpCAM⁺), and luminal committed cells (LCs: Lin⁻CD49f^low/⁻^EpCAM⁺)—from published datasets (microarray **(B)** and RNA-seq **(C)**) reveals selective enrichment of *KIT* transcripts in LPs. **(D–F)** scRNA-seq analysis of normal human breast tissue demonstrates distinct RTK expression across epithelial subsets. *cKIT* expression is largely confined to LP clusters. **(E)**, while *EGFR* is enriched in BC and LC populations **(F)**. **(G)** Representative flow cytometry profile showing CD49f and EpCAM expression in human mammary cells; cKIT⁺ cells (blue) map to the LP gate, overlaid on the total epithelial population (grey). **(H)** qRT-PCR analysis of sorted mammary subpopulations confirms that *cKIT* is primarily expressed in LPs. *SCF/KITL* is poorly expressed by mammary epithelial cell types. **(I)** Immunoblot analysis validates selective cKIT protein expression in LPs among freshly sorted BC, LP, and LC fractions from reduction mammoplasty samples. **(J)** ChIP-sequencing reveals an active chromatin landscape (H3K4me3, H3K36me3) at the *KIT* locus in LPs, contrasted by Polycomb-mediated repression (H3K27me3) in BC and LC cells. **(K)** Immunohistochemical staining identifies cKIT⁺ cells within ductal and alveolar structures of normal breast tissue, consistent with dispersed LP localization. **(L)** Mammary repopulating unit (MRU) activity show that cKIT⁺ cells isolated from the CD49f⁺ compartment exhibit reduced regeneration of colony-forming cells compared to cKIT⁻ counterparts, consistent with a lineage-restricted progenitor phenotype.

We further validated these results using FACS analysis, which confirmed surface enrichment of cKIT protein in the LP compartment-specific manner (**Figure 1G**). The quantitative-polymerase chain reaction (qPCR) and Western blot analyses of sorted populations independently corroborated the selective expression of cKIT in LP cells, with minimal or absent expression in LC and BC fractions (**Figure 1H-I**). However, the expression of the cKIT ligand (KITL)—also known as stem cell factor (SCF) or steel factor—was not found to be high in any mammary epithelial subpopulations, nor was it notably expressed by mammary stromal cells (**Figure 1H, Supplementary Figure S1B**). Given that SCF can exist as a membrane-bound protein and function through potent paracrine mechanisms, it is noteworthy to see some SCF expression in LC subsets compared to LPs (**Supplementary Figure S1B**). This suggests the possibility of paracrine cKIT signaling directed toward the LP population.

To assess the reliability of cKIT as a surface marker for mammary luminal progenitors, we first optimized its detection by immunoblotting. Only a C-terminal-directed antibody consistently recognized full-length cKIT in mammary epithelial lysates (**Figure 1I**), whereas an N-terminal antibody failed to do so (**Supplementary Figure S2A**). Single-cell suspensions from human breast tissue showed evidence of cKIT degradation compared to intact mammary tissue-organoids (**Supplementary Figure S2B**). We next evaluated epitope sensitivity to enzymatic dissociation. CD34⁺ hematopoietic stem/progenitor cell-enriched fraction retained stable cKIT surface staining, confirming that the epitope itself is not inherently labile. Time-course FACS analysis revealed a progressive loss of cKIT signals in dissociated mammary cells, with up to 32.9% reduction in the cKIT⁺ population by 3 hours post-digestion (**Supplementary Figure S2C**). These findings highlight the potential for cKIT surface loss during enzymatic dissociation, cautioning that standard single-cell workflows may underestimate the frequency of cKIT⁺ mammary epithelial cells.

To explore the chromatin landscape underlying this selective expression, we examined histone modifications at the cKIT locus using our recently published chromatin immunoprecipitation (ChIP)-sequencing method for H3K4me3, H3K27me3, and H3K36me3 (5). In LP cells, the cKIT promoter was marked by strong H3K4me3 and gene body enrichment of H3K36me3, consistent with an actively transcribed state (**Figure 1J**). In contrast, both LC and BC cells exhibited high levels of H3K27me3 and presence of H3K4me3 at the cKIT locus, indicative of a poised chromatin state (**Figure 1J**), implying that some rare non-LP epithelial cells are epigenetically primed for activation in response to appropriate developmental or signaling cues. Taken together these epigenomic signatures align with the observed expression patterns and suggest that the cKIT⁺ was not only differentially expressed, but also dynamically regulated by progenitor cell-specific chromatin states in the human mammary luminal epithelium.

To determine whether cKIT-expressing LPs occupy specific anatomical niches, we performed immunostaining on 3 normal human mammary tissues from reduction mammoplasties. cKIT⁺ LP cells and cKIT⁻ LC cells were distinctly identified across both ductal and alveolar regions, highlighting the widespread epithelial distribution of LPs within the adult human breast (**Figure 1K and Supplementary data S1C**). This spatial localization reinforces the conclusion that cKIT identifies a functionally relevant luminal population distributed throughout the mammary epithelium. These observations support the utility of cKIT as a biomarker of the luminal progenitor state within intact tissue architecture.

Next, to rule out if rare cKIT+ cells have mammary repopulating unit (MRU) activity which is restricted to BCs and to further evaluate if cKIT+ LPs contribute to stem cell activity, we sorted CD49f⁺ primitive epithelial cells prospectively into cKIT⁺ and cKIT⁻ subsets without EpCAM gating, which normally distinguishes BCs from LPs. Subsequently, the cells were embedded in collagen gels and transplanted under the renal capsule of immunodeficient mice, as described by Eirew et al (1). As expected, MRU activity was found to be restricted to cKIT-basal subset.

Following in vivo passage, cells from cKIT^+^ grafts formed fewer CFCs in ex vivo colony-forming assays compared to cKIT grafts (**Figure 1L**). This dissociation between clonal output and progenitor marker expression suggested that cKIT^+^ LPs may exist in a more luminal lineage-committed progenitor state. Together, these data established a robust, lineage-restricted expression pattern of the cKIT⁺ in human mammary epithelium, highlighting it’s potential functional relevance within the LP compartment, specifically.

### Models of LP-state Reveal cKIT-Dependent Mitogenic Signaling and Synergy and Transactivation with EGF Stimuli

To assess the functional role of cKIT signaling in mammary epithelial cells, we engineered a panel of MCF10A, estrogen receptor (ER)/progesterone receptor (PR)-negative cell lines, to express either wild-type cKIT (MCF10A-cKIT^WT^), a kinase-dead dominant-negative mutant bearing a lysine-to-methionine substitution at position 623 within the ATP-binding site of the kinase domain (MCF10A-cKIT^K623M^), or GFP as a control (MCF10A-GFP), alongside parental MCF10A cells. FACS analysis confirmed the absence of endogenous cKIT⁺ expression on parental MCF10A lines (**Figure 2A**). We next evaluated the proliferative response of these lines to graded concentrations of EGF (62.5-0 pg/mL) and SCF (100-0 ng/mL) using MTT-based proliferation assays. As expected, all lines exhibited dose-dependent growth in response to EGF (**Figure 2B**). However, only MCF10A-cKIT^WT^ cells showed a robust growth response to SCF in the absence of EGF, with proliferation reaching ∼65-85% of the EGF-induced maximum (**Figure 2B**). Cells expressing the kinase-dead mutant or controls failed to respond to SCF, underscoring the requirement for intact cKIT kinase activity in driving this effect (**Figure 2B**). Additionally, we quantified the proliferative output per input cell by counting cells for 2 consecutive passages of engineered MCF10A variants in presence of SCF. MCF10A-cKIT^WT^ cells exhibited significantly higher total cell yield per seeded cell compared to MCF10A-cKIT^K623M^, MCF10A-GFP expressing, and parental controls (**Supplementary Figure S3A**). Consistently, clonogenic assay confirmed that only the cKIT^WT^ line generated robust clonogenic output in response to SCF, with both colony number and size significantly greater than in other lines (**Supplementary Figure S3B**). The expression of oncogenic cKIT variants cKIT^D816Y^ and cKIT^D816V^ did not enhance proliferation or clonogenic capacity in MCF10A cells. Moreover, when co-expressed with cKIT^WT^ or other variants in various combinations using a dual transduction strategy (puromycin selection followed by GFP sorting), these constitutively active mutants reduced cellular fitness compared to cells expressing cKIT^WT^ alone, suggesting that constitutive activation of cKIT is not tolerated in this context (**Supplementary Figure S3A-B**).

**Figure 2.**
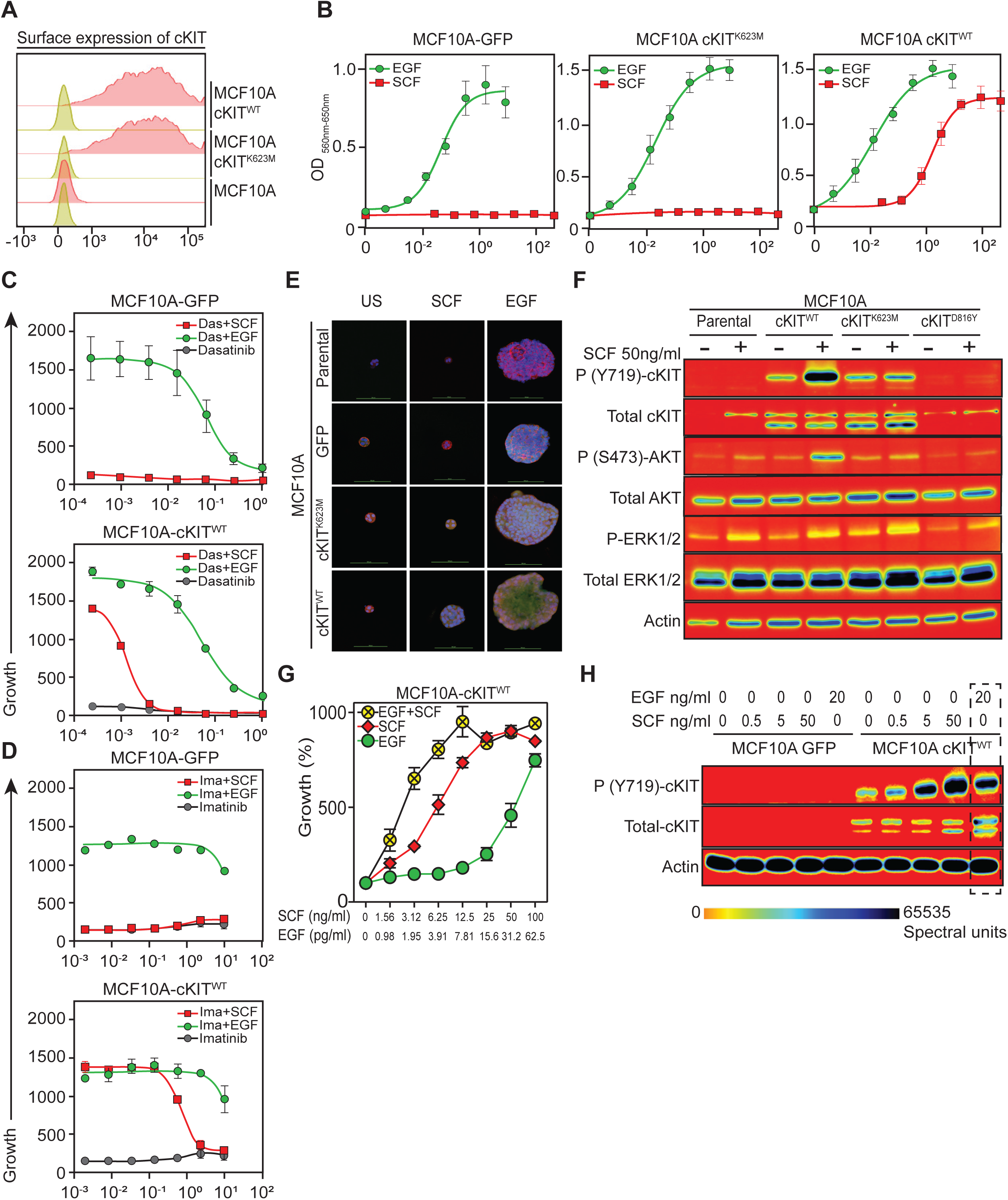
Engineered LP-State Models Reveal cKIT-Dependent Mitogenic Signaling, Synergy, and Transactivation with EGF. **(A)** Flow cytometry shows that parental MCF10A cells lack endogenous cKIT expression, while engineered MCF10A lines expressing either wild-type cKIT (cKIT^WT^) or kinase-dead mutant cKIT (cKIT^K623M^) display robust surface cKIT staining. **(B)** MTT proliferation assays demonstrate that only cKIT^WT^ cells respond to SCF in a dose-dependent manner. Neither cKIT^K623M^ nor GFP control lines show a response. All lines remain responsive to EGF. **(C–D)** SCF-driven proliferation in cKIT^WT^ cells is blocked by the tyrosine kinase inhibitors imatinib and dasatinib, confirming cKIT-specific signaling. **(E)** In 3D Matrigel assays, only cKIT^WT^ cells form enlarged organoids in response to SCF. In contrast, control lines fail to respond to SCF but grow in the presence of EGF, indicating that SCF-dependent growth is specific to cKIT activity. **(F)** Western blot analysis shows SCF-induced phosphorylation of cKIT (Y719) and downstream AKT (S473) selectively in cKIT^WT^ cells. Phosphorylation of ERK1/2 is not markedly changed. **(G)** Combined treatment with low doses of SCF and EGF produces a synergistic increase in proliferation in cKIT^WT^ cells, suggesting cooperativity between the two RTK pathways. **(H)** Western blotting reveals that EGF alone induces low-level cKIT phosphorylation in cKIT^WT^ cells, suggesting ligand-independent transactivation of cKIT by EGFR.

Moreover, morphological differences were observed between EGF- and SCF-treated MCF10A-cKIT^WT^ cells seeded at similar densities in EGF-free media, with SCF-treated cells displaying a more clustered appearance suggesting that SCF-cKIT signaling may promote cell–cell adhesion, coordinated proliferation, or epithelial organization, features that could be important during early luminal epithelial development (**Supplementary Figure S4**). Further, to confirm pathway specificity, we treated MCF10A-cKIT^WT^ cells with SCF (50 ng/mL) in the presence of tyrosine kinase inhibitors: imatinib or dasatinib. Both agents significantly attenuated SCF-induced proliferation, demonstrating that the observed response was cKIT-dependent (**Figure 2C-D**).

To investigate the functional consequences of cKIT activation in a more physiological context, we performed 3D organoid cultures and live imaging of engineered MCF10A lines in the presence or absence of SCF or EGF. In the absence of EGF, only MCF10A-cKIT^WT^ organoids exhibited increase in size, and cellularity upon SCF stimulation, whereas MCF10A-cKIT^K623M^ and MCF10A-GFP organoids showed no growth response to SCF and were similar to unstimulated controls. However, when cultured in EGF-containing medium, all lines formed similarly sized organoids with comparable morphology and size, indicating that the differential response was specific to SCF-mediated activation of cKIT (**Figure 2E**). These results suggest a functional role for ligand-dependent cKIT signaling in driving mammary epithelial progenitor expansion in MCF10A model.

To understand downstream signaling mechanisms underlying this phenotype, we performed Western blot analysis as expected, we only observed cKIT protein expression in the engineered MCF10A lines, with no detectable cKIT in the parental cells, even after SCF (50 ng/ml) stimulation. However, the expression was more robust in MCF10A-cKIT^WT^ and MCF10A-cKIT^K623M^ lines. Nevertheless, only MCF10A-cKIT^WT^ exhibited robust phosphorylation of cKIT at Y719 upon SCF stimulation. Furthermore, SCF treatment also led to strong phosphorylation of AKT at S473 in the MCF10A-cKIT^WT^ line, while total AKT levels remained unchanged across all lines. In contrast, there were no appreciable changes in phospho-ERK1/2 or total ERK levels across any condition (**Figure 2F**). cKIT variants, although established oncogenes in other tissues, exhibited modest cKIT activation when expressed alone or in combination, but failed to activate downstream AKT signaling—indicating an absence of pro-growth, constitutive kinase signaling in this context. (**Supplementary Figure S3C**). These results suggest that ligand-induced activation of cKIT selectively engages the PI3K-AKT axis in MCF10A-cKIT^WT^ cells, providing a mechanistic basis for the observed proliferative advantage and reinforcing the specificity of the cKIT-dependent mitogenic program.

We further explored any potential synergy between cKIT and EGFR signaling, towards this we co-treated MCF10A-cKIT^WT^ cells with SCF and EGF at various concentrations in MTT assays. The most striking cooperative effect was observed at subthreshold concentrations of SCF (12.5 ng/mL) and EGF (7.81 pg/mL), where combined treatment resulted in a 2- to 5-fold increase in proliferation compared to either ligand alone at the same doses (**Figure 2G**). It is also worth noting that we observed a SCF dose-dependent increase in phospho-cKIT (Y719) in MCF10A-cKIT^WT^ cells (**Figure 2H**), confirming effective ligand-dependent receptor activation. We also detected phospho-cKIT (Y719) in MCF10A-cKIT^WT^ cells treated with EGF alone, suggesting transactivation between EGFR and cKIT pathways in proliferative MCF10A luminal epithelial model.

Taken together, we demonstrate that cKIT activation is sufficient to drive a potent mitogenic program in mammary epithelial cells, which could be further amplified by low-level EGFR stimulation, highlighting a synergistic axis with implications for LP-state maintenance.

### H₂O₂-Dependent RTK Activation in Luminal Progenitor Cells: A ‘Floodgate’ Signaling Model

Transient EGFR activation has been shown to disrupt redox homeostasis by inactivating peroxiredoxins (PRDXs) and phosphatases within lipid raft-associated signaling hubs, tipping the balance toward sustained kinase activity, a process termed “floodgate” oxidative signaling (9). We previously reported elevated expression of multiple PRDXs in luminal epithelial cells compared to basal cells, suggesting a lineage-specific redox buffering capacity (3). Given our observation that EGF stimulation leads to cKIT phosphorylation, we hypothesized that cKIT may function as a redox-responsive receptor. To test this, we titrated hydrogen peroxide (H₂O₂) across engineered MCF10A models to assess cKIT activation under oxidative stress. MCF10A-cKIT^WT^ cells exhibited a dose-dependent increase in growth in response to H₂O₂ compared to MCF10A-cKIT^K623M^, and MCF10A-GFP, indicating that redox-driven proliferation requires functional cKIT kinase activity (**Figure 3A**). We further evaluated the proliferative impact of H_2_O_2_ signaling by co-stimulating MCF10A-cKIT^WT^ cells with SCF or EGF and H₂O₂. Combined treatment produced greater proliferative output than either stimulus alone (**Figure 3B**), indicating that redox and ligand inputs can act synergistically to amplify mitogenic cKIT signaling.

**Figure 3.**
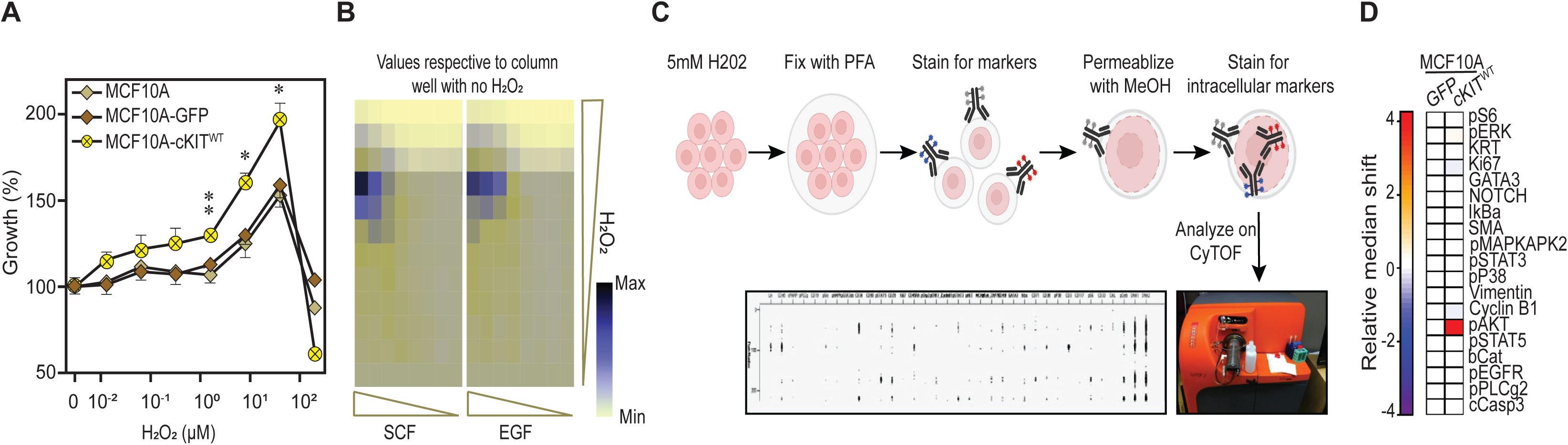
Redox-Mediated Activation of cKIT Drives Mitogenic Signaling in Luminal Progenitor-Like Cells. **(A)** MTT proliferation assays show that H₂O₂ induces a dose-dependent increase in proliferation of MCF10A–cKIT^WT^ cells. In contrast, cells expressing kinase-dead cKIT^K623M^ or GFP controls display minimal response, indicating that redox-driven proliferation requires intact cKIT kinase activity. **(B)** Co-treatment with H₂O₂ and either SCF or EGF leads to a significantly greater proliferative response in MCF10A–cKIT^WT^ cells than either stimulus alone, suggesting that oxidative and ligand-mediated inputs synergize to enhance cKIT-driven signaling. **(C–D)** Mass cytometry (CyTOF) analysis shows that exposure to 5 mM H₂O₂ selectively induces phosphorylation of AKT in cKIT^WT^-expressing cells, while vehicle-treated controls show no activation.

Further, we performed mass cytometry (CyTOF) analysis on MCF10A-cKIT^WT^ and MCF10A-GFP cells cultured in the presence or absence of H₂O₂. H₂O₂ treatment induced robust phosphorylation of AKT in MCF10A-cKIT^WT^ cells, while MCF10A-GFP controls showed minimal activation, underscoring the specificity of the redox-sensitive proliferative response (**Figure 3C**). These results highlight a novel role for cKIT as a redox-sensitive mitogenic receptor that can integrate environmental oxidative cues with growth factor signaling to enhance epithelial proliferation.

To assess whether oxidative stress could activate endogenous cKIT signaling in primary mammary epithelium, we focused on the LP subpopulation, which expresses cKIT and resistant to reactive oxygen species (ROS) (3). Sorted LP cells were embedded in 3D Matrigel domes and cultured with or without H₂O₂ for 10-12 days (**Figure 4A**). Organoids derived from H₂O₂-treated LP cells appeared consistently larger than untreated controls (**Figure 4B**). To quantify these changes, we developed a custom imaging workflow combining two-photon fluorescence and reflectance confocal microscopy to measure both total organoid structure and lumen diameters (**Figure 4C-D**). Quantitative analysis across LP derived organoids from two donors (*N=2*) showed a marked increase in both structural and luminal diameters in H₂O₂-treated organoids (**Figure 4E**), supporting the notion that oxidative stress promotes organoid growth, potentially through activation of endogenous RTK signaling.

**Figure 4.**
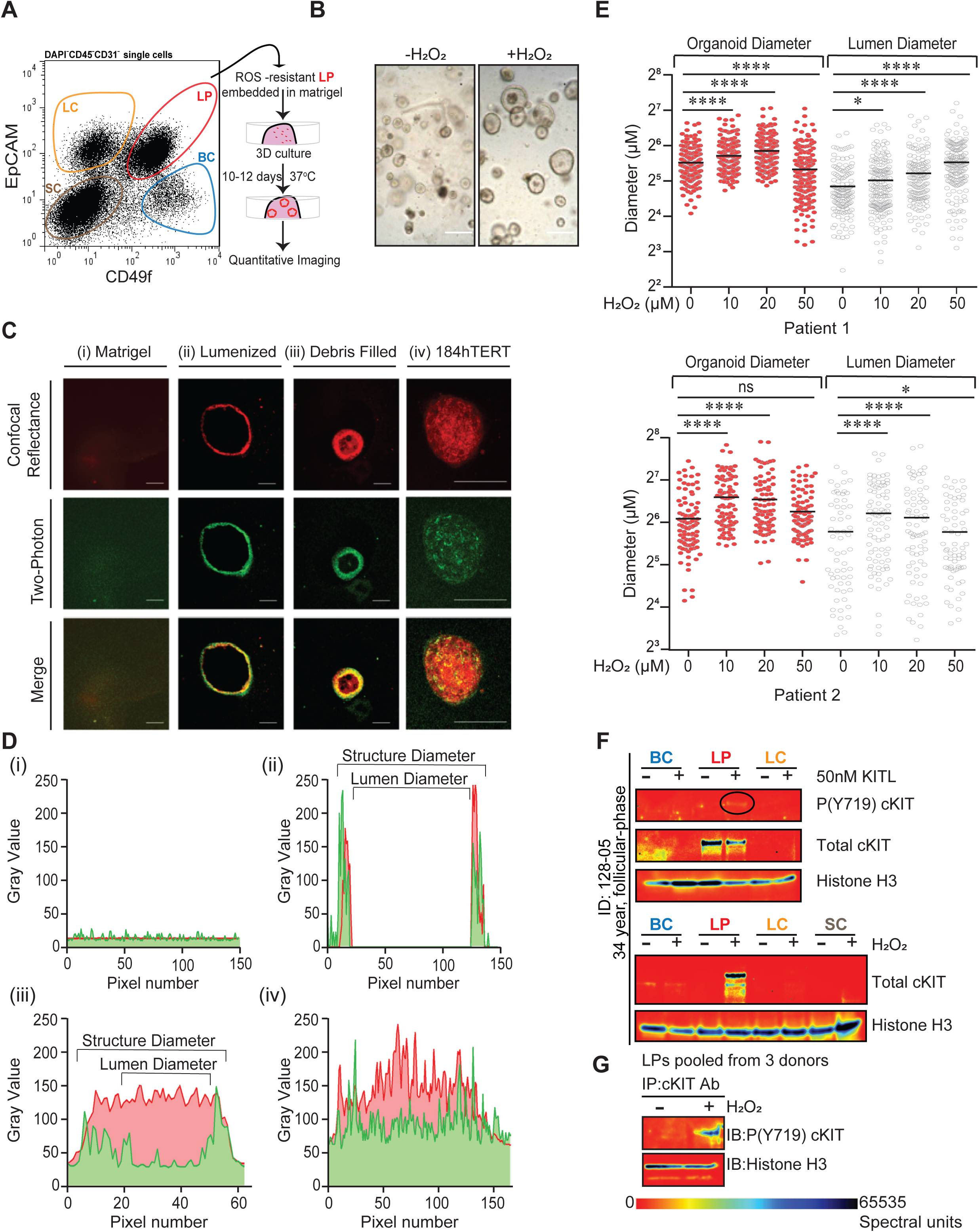
Oxidative Stress-associated cKIT Activation and Growth in LP cells. **(A)** Schematic overview of experimental workflow: FACS-purified LP cells from human mammary tissue were embedded in 3D Matrigel and cultured with or without H₂O₂ for 14 days. **(B)** Representative brightfield images reveal that H₂O₂-treated LP-derived organoids exhibit increased overall size compared to untreated controls, indicating enhanced proliferative capacity. **(C)** Two-photon fluorescence and reflectance confocal microscopy of 14-day-old LP-derived 3D structures demonstrate the formation of organized, lumenized acini. Panels include: (i) Matrigel-only background, (ii) LP-derived acinus with clear lumen and no debris, (iii) LP-derived acinus containing intraluminal debris, and (iv) disorganized multicellular 3D structure from 184-hTERT cells lacking a defined lumen. Co-registered signals from both imaging modalities appear yellow in merged images. Scale bars: 50 μm. **(D)** Pixel intensity plots along single-pixel cross-sections illustrate signal distribution within individual structures. LP-derived acini show reduced two-photon signal in the luminal space, confirming lumen formation, while 184-hTERT structures lack clear luminal architecture. **(E)** Quantification of total structure and lumen size from two independent donor-derived LP organoid cultures (*N*=2 donors) reveals significant enlargement upon H₂O₂ treatment, consistent with redox-stimulated growth. **(F)** Western blot of FACS-purified basal (BC), luminal progenitor (LP), and luminal-committed (LC) cells treated with SCF (50 ng/mL) shows cKIT phosphorylation at Y719 selectively in LPs. **(G)** Immunoprecipitation–Western blot (IP-WB) of pooled primary LPs from three independent donors confirms that H₂O₂ exposure induces robust phosphorylation of cKIT at Y719, demonstrating redox-mediated activation of endogenous cKIT.

Following this, we assessed whether oxidative stress could activate endogenous cKIT signaling in primary mammary LP subpopulation. We purified BC, LP, and LC cells from three reduction mammoplasty specimens and performed Western blotting after treatment with 50 ng/ml SCF or 100µM H₂O_2_. Phosphorylation of cKIT (Y719) was observed exclusively in LPs following either SCF or H₂O₂ stimulation, with no detectable expression in BC or LC fractions (**Figure 4F**). To validate these findings, we purified LPs from 3 reduction mammoplasty specimens and pooled LPs from all three donors and performed immunoprecipitation using cKIT antibody and immunoblotting using Y719 phosphorylation specific antibody (IP-WB), which confirmed robust cKIT activation in response to H₂O₂ stimuli (**Figure 4G, Supplementary Figure S5A**). Notably, despite the absence of serum, H₂O₂ alone, was sufficient to induce extensive global phospho-tyrosine signaling in mammary epithelial cells (**Supplementary Figure S5B**). Further analysis revealed that luminal epithelium exhibited elevated H₂O₂-induced protein tyrosine residue phosphorylation relative to basal lineage and stromal cell (SC) populations (**Supplementary Figure S5C**). These data position LPs as a dominant redox-sensitive subpopulation in the human mammary epithelium, with cKIT acting as a key effector. Previously, we reported that LPs exhibit unique metabolic profile characterized by glutathione independence and enhanced resistance to H₂O₂ induced oxidative stress distinguishing them from basal myoepithelial lineage which rely on glutathione-dependent antioxidant mechanisms (3). To determine whether H₂O₂ responsiveness extended beyond cKIT activation, we examined the phosphorylation status of peroxiredoxin 1 (PRDX1), a key redox sensor and signaling relay enzyme. We first established H₂O₂-dependent PRDX1 inactivation in HEL cells (13), observing a dose-dependent increase in phospho-PRDX1 (**Figure 5A**). HEL cells endogenously express high levels of cKIT and therefore provide a well-characterized system to study redox regulated signaling. Further, our analysis shows that even in the absence of fetal bovine serum, H₂O₂ alone is sufficient to induce PRDX1 phosphorylation in HEL cells and bulk mammary single cells (**Figure 5B**). We then sorted single mammary cells into three epithelial subsets BC, LP, and LC as well as a stromal (SC) population from three independent reduction mammoplasty donors (*N*=3 donors). Following H₂O₂ stimulation, phospho-PRDX1 levels were elevated predominantly in luminal lineage cells, with LPs showing the highest signal. Notably, LPs also exhibited constitutively elevated phospho-PRDX1 levels in the absence of exogenous H₂O₂, suggesting a chronic redox activation state distinct from both BC and LC populations (**Figure 5C**). This constitutive PRDX1 inactivation across LPs from donors differing in hormonal (luteal and follicular) status supports the notion that LPs maintain an intrinsically redox-primed state. To explore whether PRDX1 is functionally associated with cKIT signaling, we performed immunoprecipitation (IP) using a cKIT-specific antibody in HEL cells and in primary LPs, followed by Western blotting. These analyses revealed co-precipitation of PRDX1 with cKIT in both systems (**Figure 5D-E**), indicating physical and potentially functional interaction. Together with our findings on redox-activated cKIT signaling, these results suggest that LPs function as specialized redox-sensing element within the mammary epithelium, poised to integrate oxidative cues into proliferative responses.

**Figure 5.**
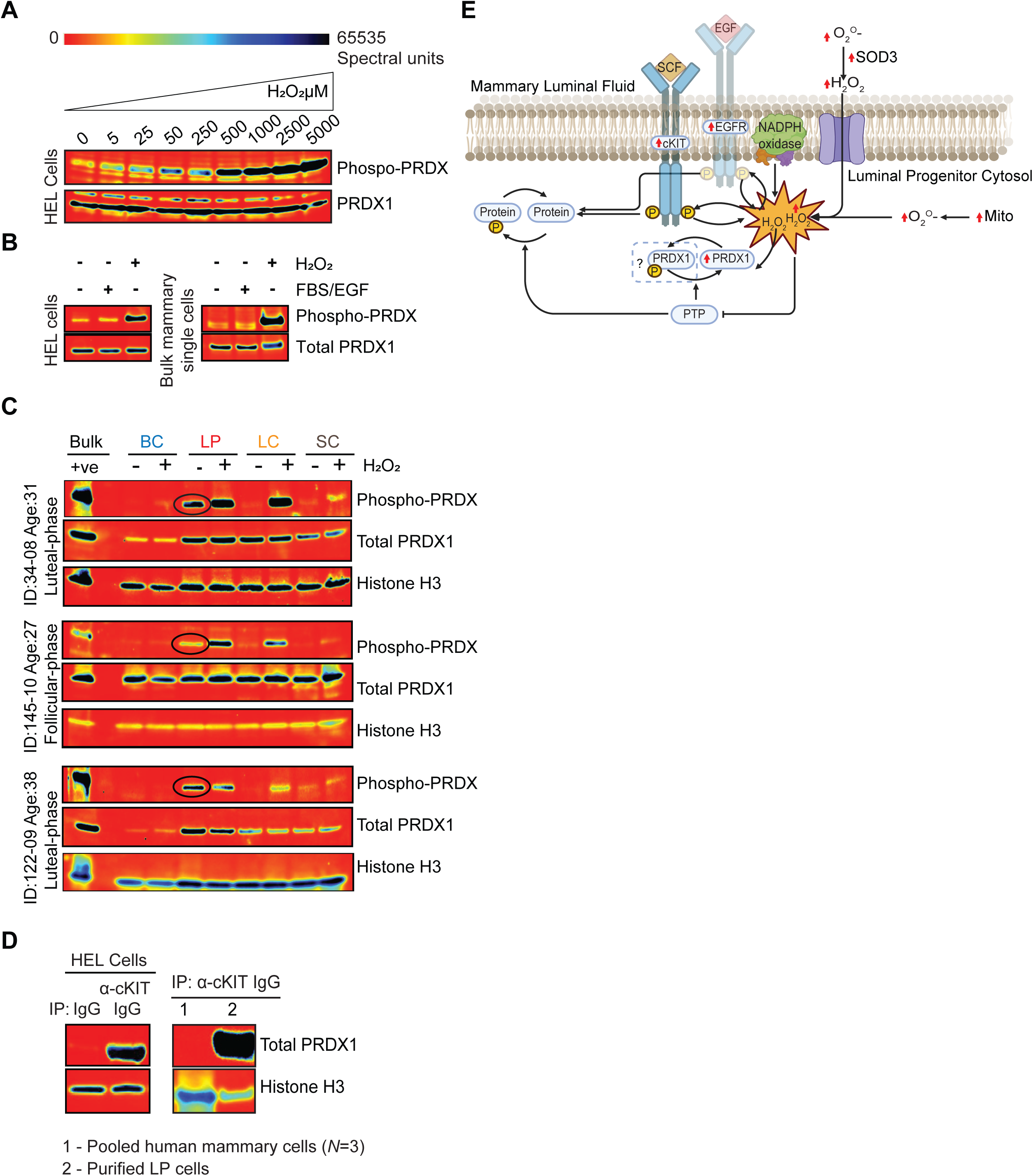
Luminal Progenitor Cells Exhibit Constitutive PRDX1 Inactivation via Phosphorylation, Reflecting an Intrinsic Redox-Primed State. **(A)** In HEL cells, increasing concentrations of H₂O₂ induce a dose-dependent rise in phospho-PRDX1 levels, confirming PRDX1 as a redox-responsive sensor of oxidative stress. **(B)** Western blot analysis shows that H₂O₂ treatment alone—without the presence of fetal bovine serum—is sufficient to induce PRDX1 phosphorylation, indicating direct oxidative inactivation independent of serum-derived growth factors. **(C)** Immunoblotting of FACS-purified BC, LP, and LC cells from three independent human mammoplasty samples reveals elevated basal levels of phospho-PRDX1 specifically in LPs. This constitutive phosphorylation pattern was consistent across donors and menstrual cycle stages (luteal and follicular), suggesting that LPs maintain a lineage-specific, redox-activated antioxidant state. **(D)** Immunoprecipitation–Western blot of HEL cells and pooled, unseparated human mammary epithelial cells and sorted LPs demonstrates that total PRDX1 is enriched in LP populations, further supporting their heightened antioxidant capacity. **(E)** Illustration showing a model integrating of SCF/EGF and redox H₂O₂ signals in human mammary luminal progenitor cells. Synergistic expression of cKIT and EGFR in LPs leads to receptor activation by their ligands (SCF and EGF), stimulates downstream signaling and promotes NADPH oxidase–mediated production of extracellular superoxide. Superoxide is converted to H₂O₂ by superoxide dismutase 3 (SOD3) and enters the cytosol. Mitochondrial respiration may also contribute to intracellular superoxide levels. Accumulation of H₂O₂ inactivates protein tyrosine phosphatases (PTPs) and leads to phosphorylation of PRDX1, which becomes functionally impaired under sustained oxidative stress. This creates a redox-positive feedback loop that amplifies RTK signaling in LPs. Solid arrows indicate known or proposed signaling events; dashed box and question mark indicate unresolved mechanisms.

### Clinical Association of cKIT Expression in Basal-Like Breast Cancer

Although cKIT expression without accompanying activating mutation has been previously reported in basal-like breast tumors using immunohistochemistry (14,15), it was not thought to influence prognosis. Given the proposed role of cKIT⁺ LPs in basal-like tumor origin (2), we re-evaluated the prognostic significance of epithelial cKIT expression in breast cancer. In addition to luminal progenitors (LPs), cKIT is also highly expressed in mast cells within the mammary gland (**Figure 6A, Supplementary Figure S6**). As mast cells represent a rare but cKIT-rich immune population, their presence could confound interpretation of epithelial-specific cKIT expression. We applied a computational regression approach to remove mast cell associated transcripts (16). Based on the bulk RNA-seq data from The Cancer Genome Atlas (TCGA) (17) PAM50 subtypes cKIT was enriched in basal-like tumors, consistent with prior studies (**Figure 6B, Supplementary Figure S7A**). Notably, high cKIT expression was significantly associated with shorter progression-free intervals (PFI), both before and after mast cell regression (**Figure 6C**). This trend was independently observed in the Molecular Taxonomy of Breast Cancer International Consortium (METABRIC)(18) dataset for survival though it did not reach statistical significance in this dataset (**Supplementary Figure S7B-C**).

**Figure 6.**
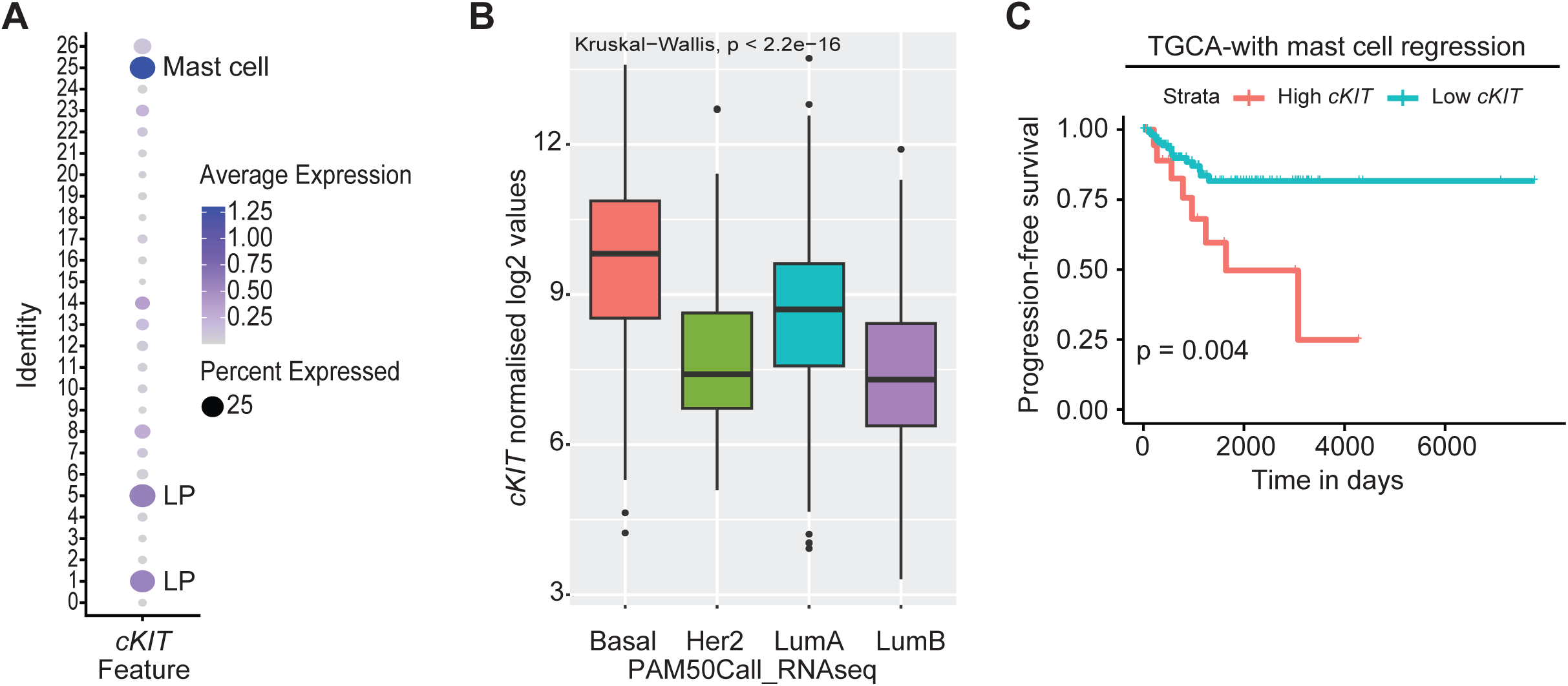
Clinical Association of cKIT Expression with Poor Prognosis in Basal-Like Breast Cancer. **(A)** Dot plot showing expression of the *cKIT* gene across 27 cell clusters identified from single-cell RNA sequencing of human breast tissue (pooled from three patients). Dot size represents the percentage of cells within each cluster expressing *cKIT*, while color intensity indicates an average expression level. *cKIT* expression is highest in mast cells (Cluster 25) and selectively enriched in two LP clusters (Clusters 1 and 5), suggesting lineage-specific transcriptional activity. **(B)** Analysis of TCGA breast cancer samples shows that cKIT expression is enriched in the basal-like molecular subtype compared to other PAM50 subtypes. **(C)** Kaplan–Meier survival analysis using progression-free interval (PFI) data from the TCGA cohort demonstrates that high cKIT expression is significantly associated with reduced survival in patients with basal-like breast cancer. To improve specificity for epithelial cKIT expression, mast cell–derived signals were computationally removed prior to analysis. Expression cutoffs for stratification were determined using the *surv_cutpoint* algorithm, which identifies optimal thresholds for prognostic discrimination rather than relying on median expression levels. P-values indicate statistical significance of survival differences.

These findings prompted us to evaluate whether cKIT activation has functional consequences in established breast cancer models. While cKIT conferred a proliferative and drug-resistant phenotype in non-transformed MCF10A cells, enforced expression of wild-type cKIT, its kinase-dead mutant, or oncogenic variants in breast cancer cell lines (T47D, MDA-MB-231, MCF7) did not affect sensitivity to RTK inhibitors (imatinib, dasatinib, dovitinib) or 5-fluorouracil across a 72-hour dose range (10⁻³ to 10¹ μM; **Supplementary Figure S8**). Growth factor and RTK dependence is a well-documented feature of many breast cancers and suggests that the cKIT signaling observed in luminal progenitors similarly depends on a lineage-specific epigenetic and proteomic context that is not preserved in transformed cell lines.

The association between high epithelial cKIT expression and poor prognosis in a subset of basal-like tumors may therefore reflect retention of a luminal progenitor-like, redox-primed state rather than canonical oncogenic cKIT activity. Consistent with this interpretation, expression of constitutively active cKIT variants reduced cellular fitness in MCF10A models, suggesting that tight regulation of cKIT levels and activation is required for maintaining the proliferative advantage of this progenitor state. Taken together, these findings establish epithelial cKIT as a candidate biomarker of a progenitor-like, stress-adapted cell state that may contribute to disease progression in basal-like breast cancer (**Supplementary Figure S8**).

Further to evaluate lineage specific redox sensitivity within the human mammary epithelium, we exposed 2D-culture expanded primary basal and luminal cells purified using two different FACS strategies (3) from two prophylactic mastectomy specimens (both from BRCA mutants) to oxidative stress induced by buthionine sulfoximine (BSO). BSO is a selective inhibitor of glutamate–cysteine ligase (GCLC), the rate-limiting enzyme in glutathione biosynthesis. In both patients, basal cells displayed a marked, dose-dependent decrease in viability following 72-hour BSO treatment, while luminal cells remained comparatively resistant (**Supplementary Figure S9A-B**), as reported previously for freshly isolated LP cells from breast tissues (3). These results further confirm that redox buffering capacity starkly differs between basal and luminal lineages. Taken together, these findings support a model in which luminal cells, particularly the cKIT^+^ LP subset, maintain enhanced antioxidant defenses that contribute to their survival under oxidative stress conditions.

## DISCUSSION

This study defines a novel redox-integrated mitogenic signaling program that is specific for luminal progenitor (LP) cells within the human mammary epithelium. Building on our previous work that showed LPs possess a glutathione-independent oxidative stress control mechanism (3), we now show that this antioxidant environment is coupled to a redox-sensitive receptor tyrosine kinase (RTK) signaling axis involving cKIT. In contrast, basal cells (BCs) rely more heavily on glutathione-dependent enzymes such as glutathione peroxidase-2 (GPX2) and exhibit lower tolerance to oxidative stress (3). In our study, we demonstrate that cKIT is selectively expressed in the CD49f⁺ LP compartment, where it is maintained in an epigenetically accessible state. In contrast, cKIT is silenced in both basal and luminal-committed (LC) cells through Polycomb-associated H3K27me3 (5). This pattern of expression marks cKIT as a feature of a transient, proliferation-competent progenitor state.

Functionally, using an engineered MCF10A model that recapitulates the luminal progenitor-like EGFR⁺cKIT⁺ receptor tyrosine kinase profile, we show that cKIT can integrate multiple inputs, its canonical ligand SCF, transactivation via EGFR, and oxidative stimuli such as H₂O₂. These cues converge on phosphorylation of cKIT and downstream activation of the AKT pathway. Importantly, LPs show constitutive phosphorylation of the antioxidant enzyme PRDX1, a peroxiredoxin known to regulate redox signaling. This supports the presence of a tonic oxidative state that permits reversible inactivation of peroxidases. This is consistent with the ‘floodgate’ model, in which oxidative conditions increase RTK activity by transiently inactivating antioxidant regulators and phosphatases, thereby amplifying downstream signaling.

This redox-tuned signaling appears to be well-adapted to the functional needs of LPs. In particular, lactation is a physiological state known to generate high levels of oxidative stress in the mammary gland (6). The ability of LPs to maintain proliferative capacity under such conditions, in part through PRDX1 inactivation and cKIT activation, likely supports rapid epithelial remodeling and secretory function during lactation. These findings offer a mechanistic basis for redox resilience in LPs and raise the possibility that similar programs may be co-opted during early stages of breast oncogenesis. Our functional assays confirm that MRU activity remains confined to cKIT⁻ basal cells. In contrast, cKIT⁺ LPs exhibit robust short-term proliferation but lack long-term regenerative capacity. This supports a developmental model in which cKIT marks a committed progenitor downstream of multipotent basal stem cells.

Clinically, we found that cKIT expression in breast tumors, after correcting for mast cell– derived signals, is associated with worse progression-free survival in both TCGA and METABRIC datasets. While cKIT is frequently silenced early during tumorigenesis through hypermethylation of promoter region (19), its retention in a subset of tumors may reflect preservation of LP-like features, including redox tolerance and proliferative responsiveness. These features could contribute to therapeutic resistance or disease progression.

In summary, this study identifies cKIT as a redox-sensitive RTK epigenetically restricted to the LP compartment of the human mammary epithelium. Our data defines a novel signaling paradigm in which oxidative cues modulate kinase activity to specifically drive LP proliferation under physiological oxidative stress. These findings advance our understanding of epithelial progenitor cell biology and provide a foundation for investigating redox-integrated RTK signaling in normal tissue function and disease.

## MATERIALS AND METHODS

### Human Mammary Tissue Procurement and Cell Sorting

Normal human mammary epithelial tissues were obtained multiple sources under institution-specific protocols: samples from the BC Cancer Research Centre (Vancouver) were collected following the protocol established by the late Dr. Connie Eaves; additional samples were acquired from Mayo Clinic Rochester under Dr. Kannan’s protocol, and from Mayo Clinic Jacksonville under Dr. Radisky’s protocol. All procedures were conducted in compliance with the Institutional Review Board (IRB) guidelines and ethical regulations at each respective institution. Upon receipt, tissue samples were mechanically and enzymatically dissociated using collagenase/hyaluronidase (Stem Cell Technologies) followed by sequential digestion with 0.25% trypsin (Stem Cell Technologies), dispase (Stem Cell Technologies), and DNase I (Sigma) to generate single-cell suspensions. Epithelial cells were subsequently stained with fluorophore-conjugated antibodies against EpCAM, CD49f, cKIT, CD45, CD31, CD34, and viability dye DAPI. FACS was subsequently performed on a FACSVantageSE, FACSDiva or a FACS Melody (Becton Dickinson). Purity of each sorted population was confirmed by FACS (1,3).

### Cell Culture and Genetic Manipulation

MCF10A cells were cultured in DMEM/F12 (Gibco) supplemented with 5% horse serum (Gibco), 20 ng/mL EGF (Sigma), 0.5 µg/mL hydrocortisone (Sigma), 10 µg/mL insulin (Sigma), 100 ng/mL cholera toxin and 1X penicillin-streptomycin (Sigma). MDA-MB-231, T47D and MCF7 lines were cultured in DMEM high glucose basal media (Gibco) supplemented with 10% fetal bovine serum and 1X penicillin-streptomycin.

### Construction of cKIT lentivectors

The *cKIT* cDNA in the pMPG(AP)-PGK-EGFP vector was generously provided by late Dr. Eaves (BC Cancer Research Institute). This cDNA was PCR-amplified and subcloned into the p3xFlag-CMV-14 expression vector (Sigma). Site-directed mutagenesis was performed using the Takara Advantage GC-Rich PCR Kit (Clontech-Takara) to generate the kinase-dead (K623M) and constitutively active (D816Y and D816V) cKIT mutants, following standard protocols. The wild-type and mutant cKIT sequences were then PCR-amplified using Q5 High-Fidelity DNA Polymerase (New England Biolabs) and subcloned into the pLMP-PGK-EGFP lentiviral vector, a modified version of pMPG(AP)-PGK-EGFP with an expanded multiple cloning site. All final constructs were sequence-verified prior to downstream applications.

### Lentiviral Production and Transduction of MCF10A Cells

High-titer lentiviral supernatants encoding human cKIT wild-type (WT), kinase-dead mutant (K623M), constitutively active mutant (D816Y), or GFP control were generated by transient transfection of HEK293T cells using a third-generation lentiviral packaging system as previously described (20). For transduction, MCF10A cells were plated at ∼50– 60% confluency and incubated with lentiviral supernatants in the presence of polybrene (8 µg/mL). After 24 hours, the media was replaced with a fresh growth medium. Forty-eight hours later, GFP⁺ cells were sorted using FACS or selected with puromycin. Subsequently, the cells were expanded for assays.

### Proliferation Assays

For MTT assays, transduced MCF10A, MDA-MB-231, MCF7 or T47D cells were seeded in 96-well plates (2,000 cells/well) and treated with SCF, EGF, hydrogen peroxide or drugs (imatinib and dasatinib). After 72 h, 10µl of MTT reagent (5 mg/mL) was added for 4 h, followed by overnight solubilization and absorbance measurement at 570 and 660nm. For direct cell counting, cells were seeded in 10 cm^2^ plates and counted using a hemocytometer after 5–6 days.

### Quantitative PCR and Western Blotting

Total RNA was extracted using Trizol (Qiagen), and cDNA was synthesized using SuperScript III (Thermo Fisher). qPCR was performed using SYBR Green Master Mix (Qiagen) on a QuantStudio 6 Flex instrument. GAPDH served as housekeeping control. For Western blotting, whole-cell lysates were prepared in RIPA buffer with protease and phosphatase inhibitors. Proteins were separated by SDS-PAGE and transferred to PVDF membranes. Membranes were incubated with antibodies against total and phosphorylated cKIT, AKT, ERK1/2, and Actin (loading control). Signal was detected using HRP-conjugated secondary antibodies and enhanced chemiluminescence (ECL) (3,21).

### Sub-renal Xenotransplantation Assay

NOD/SCID mice were anesthetized, the dorsal fur removed, and the skin disinfected with 70% ethanol. A ∼1.5 cm midline incision was made to expose the kidneys, and a small cut in the abdominal wall allowed each kidney to be gently externalized. Using a dissecting microscope, a 2–4 mm slit was made in the renal capsule, and up to four collagen gels were inserted beneath using a fire-polished pipette. The abdominal wall was closed with sutures, and the same procedure was performed on the opposite kidney. To support hormonal conditions, a subcutaneous slow-release pellet (2 mg β-estradiol, 4 mg progesterone in MED-4011 silicone) was implanted at a separate site before closure of skin. This regimen produces mid-luteal phase–like hormonal levels in experimental animals (22).

For recovery of regenerated cells, mice were euthanized, and collagen gels were dissected under sterile conditions. Gels were digested at 37 °C for 4–4.5 hours in SF7 medium with 5% FBS, collagenase (600 U/mL), and hyaluronidase (200 U/mL), followed by trypsin-EDTA treatment and gentle pipetting. All animal procedures were approved by the UBC Animal Care Committee (A06-1520) (1,3).

### In Vitro Colony-Forming Cell (CFC) Assay

CFC assays were performed by seeding mammary epithelial cells obtained from primary tissues or dissociated collagen gels onto 60-mm culture dishes pre-coated with a 1:43 dilution of Vitrogen 100 collagen (Collagen Biotechnologies, Palo Alto, CA) in PBS. Each dish received 2.0 × 10⁵ irradiated (50 Gy) NIH 3T3 fibroblast feeder cells in 4 mL of EpiCult-B medium (Stemcell Technologies) supplemented with 5% FBS and 0.5 µg/mL hydrocortisone. After 24 hours, cultures were switched to serum-free EpiCult-B medium with hydrocortisone or to SF7 medium (DMEM/F12 supplemented with 0.1% BSA, 10 ng/mL EGF, 10 ng/mL cholera toxin, and 1 µg/mL insulin). After 7–10 days of incubation at 37 °C and 5% CO₂, colonies were fixed with a 1:1 methanol: acetone solution and stained with Wright’s Giemsa. Colonies were scored under a dissecting microscope and categorized into luminal, myoepithelial, or mixed lineages based on morphology (1).

### Single Cell Mass Cytometry (CyTOF)

MCF10A-cKIT^WT^ and GFP cells were harvested using trypsin and treated with 5mM H_2_O_2_ for 10 min and pelleted by centrifugation. Subsequently, cells were washed with cell staining buffer (CSB; Fluidigm) and incubated with cisplatin (5 µM, 5 min) for viability staining. After quenching with CSB and washing, cells were fixed with 1.6% paraformaldehyde (PFA) for 10 minutes at room temperature. Cells were then labeled with a palladium-based 20-plex barcoding kit (Fluidigm) according to the manufacturer’s protocol to enable multiplexed acquisition. Following barcoding, samples were pooled and blocked with Fc receptor blocker before staining with a cocktail of metal-conjugated antibodies directed against surface and intracellular markers. Metal isotope-conjugated antibodies were either obtained from Fluidigm or labeled in-house using the Maxpar X8 labeling kit (Fluidigm). For intracellular staining, cells were permeabilized in methanol (chilled, 90% v/v) and incubated with intracellular antibodies overnight at 4°C. After washing, cells were labeled with intercalator-Ir (191/193Ir, Fluidigm) in 1.6% PFA for 20 minutes at room temperature for DNA content analysis. Samples were acquired on a Helios mass cytometer (Fluidigm), using EQ Four Element Calibration Beads for normalization. Normalized .fcs files were debarcoded using the Premessa R package. Downstream analyses were conducted in Cytobank and R, with dimensionality reduction (viSNE) and clustering (FlowSOM) used to visualize and quantify epithelial cell states and signaling pathway activity (4).

### Multi-Modal Imaging and 3D Image Analysis

Dual-modality imaging was performed using a custom-built experimental microscope described elsewhere (23) optimized for reflectance confocal and two-photon microscopy. Images were acquired concurrently at a resolution of 256 × 256 pixels, representing a 300 × 300 µm field of view, using an excitation wavelength of 720 nm for both modalities.

Following acquisition, images were analyzed using ImageJ software. For quantifying 3D mammary structure dimensions, the line measurement tool was used to measure diameters at the widest and narrowest points through the structure’s lumen, and the average of the two was used to minimize directional bias. Pseudocolor overlays, 3D reconstructions, and pixel intensity analyses were performed using ImageJ’s built-in functions (24,25). Co-localization was assessed using the Just Another Co-localization Plugin (JACoP), and Pearson correlation coefficients were calculated to quantify signal overlap. All statistical analyses were conducted using GraphPad Prism.

### Single Cell RNA Sequencing and Data Analysis

Human breast tissue was dissociated into single cells and sorted for DAPI-viable fraction and resuspended in PBS,and stained with trypan blue and counted using hemocytometer. The volume of cell suspension was adjusted to 1000 cells/ul in PBS. Cells were then processed on a 10x Genomics Chromium Controller using the Chromium Next GEM Single Cell 3′ reagents kit v3.1 (Dual Index) following the manufacturer’s instructions. In brief, approximately 16,500 live cells were loaded onto the Chromium controller to recover 10,000 cells for library preparation and sequencing. Gel beads were prepared following the manufacturer’s instructions. Subsequently, oil partitions of single-cell and oligo coated gel beads were captured, and reverse transcription was performed, resulting in cDNA tagged with a cell barcode and unique molecular index (UMI). Next, GEMs were broken, and cDNA was amplified and quantified using an Agilent Bioanalyzer High Sensitivity chip (Agilent Technologies).

To prepare the final libraries, amplified cDNA was enzymatically fragmented, end-repaired, and polyA tagged. Fragments were then selected, according to their size using SPRIselect magnetic beads (Beckman Coulter). Next, Illumina sequencing adapters were ligated to the size-selected fragments and cleaned up using SPRIselect magnetic beads (Beckman Coulter). Finally, sample indices were selected and amplified, followed by a double-sided size selection using SPRIselect magnetic beads (Beckman Coulter). Final library quality was assessed using an Agilent Bioanalyzer High Sensitivity chip. The libraries were then sequenced as paired end reads (PE150) on Novaseq platform (illumina)(12,26).

### Transcriptomic Data Analysis

Single-cell RNA sequencing (scRNA-seq) data were processed and analyzed using the Seurat v4 package in R. Raw gene expression matrices were log-normalized and scaled following quality control filtering to exclude cells with low RNA content or high mitochondrial gene expression. Dimensionality reduction was performed using principal component analysis (PCA), followed by clustering and visualization with Uniform Manifold Approximation and Projection (UMAP). Differential gene expression analysis was performed using a Wilcoxon rank-sum test, corrected for multiple comparisons using the Bonferroni method. For visualization and lineage annotation, we integrated publicly available human mammary epithelial datasets and matched RTK expression to defined cell states (basal, luminal progenitor, and luminal committed). Gene ontology and pathway enrichment analyses were conducted using the clusterProfiler and ReactomePA R packages(26).

### Clinical Dataset Analysis (TCGA and METABRIC)

Clinical and transcriptomic data for breast cancer patients were obtained from The Cancer Genome Atlas (TCGA) and Molecular Taxonomy of Breast Cancer International Consortium (METABRIC) via cBioPortal and the UCSC Xena browser. For each dataset, PAM50 molecular subtype was also obtained from clinical data sheet. Normalized mRNA expression levels of *cKIT* and other relevant genes were extracted and analyzed in R and analyzed using R (v4.3).

To assess clinical relevance, we examined associations between *cKIT* expression and progression-free survival (PFS) using Kaplan-Meier survival curves and log-rank tests. Patients were stratified into high and low *cKIT* expression groups based on upper and lower quartiles or using an optimized cutpoint derived from maximally selected rank statistics (R package survminer).

To distinguish epithelial-specific *cKIT* expression, mast cell–associated gene signatures were computationally removed or regressed out using established marker panels (e.g., *TPSAB1*, *CMA1*)(16,27,28). All statistical analyses were performed using R (v4.2) and visualized using ggplot2 and Kaplan–Meier survival curves.

## AUTHOR CONTRIBUTIONS

S.M.M.A. contributed to data generation and manuscript drafting. M.R. performed single-cell RNA sequencing analysis, while G.S. led molecular cloning. Data generation was also supported by K.S., Y.D., G.K., C.M., and Y.Y. D.K. conducted CyTOF analysis, and P.E. carried out MRU assays. Epigenomic analyses were performed by D.P., and C.P.E. provided manuscript review. H.W. and H.Z. contributed multimodal microscopy imaging. J.J. supported funding acquisition. A.D., S.ML., and J.B. provided surgical specimen access. M.G., M.S., and D.R. offered critical review and input on the manuscript. A.S. and H.P. performed TCGA and METABRIC data analysis. N.K. led the conceptualization, study design, manuscript drafting, supervision, and funding acquisition, and contributed to both data generation and analysis. All authors reviewed and approved the final manuscript.

## ACKNOWLEDGEMENTS

Dr. Connie Eaves passed away during the preparation of this manuscript. Her scientific insights and contributions to this work are gratefully acknowledged. We acknowledge members of Dr.Eaves and Dr.Kannan laboratories for technical assistance. We thank Dr. Sue Goo Rhee (Ewha Womans University, Seoul, South Korea) for generously providing the phospho-PRDX1 antibody.

## SUPPLEMENTARY DATA

**Supplementary Figure S1. cKIT is Selectively Expressed in Luminal Progenitor (LP) cells of the Human Mammary Epithelium.**

**(A)** Transcriptomic profiling of FACS-purified mammary epithelial subpopulations BC, LP, and LC cells—reveals that cKIT transcripts are enriched specifically in LPs. Data are compiled from previously published microarray (10) and RNA-seq (11) datasets.

**(B)** *SCF/KITLG* expression levels in human mammary epithelial populations from the Pellacani et al. dataset (5) highlight potential paracrine signaling within the mammary gland.

**(C)** Immunohistochemical staining of normal breast tissue sections (*N*=2 donors) confirms cKIT protein expression is confined to luminal cells with progenitor morphology in both ductal and alveolar structures.

**Supplementary Figure S2. Surface cKIT is sensitive to trypsin-mediated cleavage.**

**(A)** Western blot of total protein lysates from mammary epithelial cells using an antibody targeting the extracellular (N-terminal) domain of cKIT fails to detect intact cKIT, suggesting susceptibility to enzymatic degradation.

**(B)** Tryptic digestion assays show that cKIT is cleaved in mammary epithelial cells upon trypsin treatment, whereas full-length cKIT is preserved in CD34⁺ hematopoietic cells and organoid-enriched mammary tissue cultures.

**(C)** Flow cytometry analysis of freshly dissociated breast tissue shows a progressive decrease in surface cKIT⁺ cell frequency within the luminal compartment over a 3-hour time course following trypsin exposure, indicating potential underestimation of LPs in single-cell workflows.

**Supplementary Figure S3. Functional consequences of cKIT activation in engineered MCF10A models.**

**(A)** Serial passage of these engineered lines in SCF-supplemented media shows sustained proliferative advantage in cKIT^WT^-expressing cells, supporting functional relevance of ligand-mediated cKIT signaling.

**(B)** Clonogenic assays demonstrate that MCF10A cells expressing wild-type cKIT (cKIT^WT^), or co-expressed with other cKIT variants, form significantly more colonies in response to SCF compared to kinase-dead (cKIT^K623M^) or control GFP-expressing cells.

**(C)** Robust phosphorylation of cKIT and AKT was observed engineered MCF10A in cell lines expressing cKIT^WT^ or in combination with other variants but not in oncogenic cKIT variants in the presence of SCF/KITL, confirming receptor activation.

**Supplementary Figure S4. Distinct morphological responses to EGF and SCF in MCF10A–cKIT^WT^ cells.**

Representative phase-contrast images of MCF10A–cKIT^WT^ cells treated with EGF (20 ng/mL) or SCF (50 ng/mL) for 72 hours. EGF-treated cells appear scattered and elongated, reflecting classical MCF10A morphology. In contrast, SCF-treated cells display a more compact, clustered appearance, indicating distinct cellular outcomes despite convergent RTK signaling.

**Supplementary Figure S5. LP cells exhibit constitutive phosphorylation of PRDX1, reflecting a redox-primed state.**

**(A)** Immunoblotting of FACS-purified BC, LP, and LC cells from two independent human mammoplasty samples (follicular stage 34- and 27-year-old) reveals elevated basal levels of phospho-PRDX1 specifically in LPs. This constitutive phosphorylation pattern was consistent across donors and menstrual cycle stages (luteal and follicular), suggesting that LPs maintain a lineage-specific, redox-activated antioxidant state.

**(B)** Western blot analysis shows that treatment with H₂O₂ alone induces phospho-tyrosine (p-Tyr) levels in mammary epithelial cells, even in serum-free conditions.

**(C)** Immunoblots of sorted epithelial BC, LP, LC, and stromal (SC) cells from three donors show elevated H2O2 induced p-Tyr in luminal epithelium specifically in LPs, supporting a redox-sensitive oxidative signaling state that distinguishes luminal from basal lineage.

**Supplementary Figure S6.** UMAP projection of single-cell RNA-seq data highlighting cKIT expression in mast cells (Cluster 25), which are known to endogenously express cKIT, while cluster 1 and 5 represent LPs.

**Supplementary Figure S7. High cKIT expression predicts poor prognosis in basal-like breast cancer.**

**(A–B)** Kaplan–Meier survival analysis using progression-free interval (PFI) data from TCGA and METABRIC cohorts reveals that high cKIT expression is associated with shorter survival in basal-like breast cancer.

**(C)** After computational regression of mast cell–derived cKIT signals, epithelial-specific cKIT expression remains significantly associated with poor prognosis. Optimal thresholds for stratification were determined using the *surv_cutpoint* method rather than median-based dichotomization. P-values reflect statistical significance.

**Supplementary Figure S8. Effect of cKIT expression on drug sensitivity in breast cancer cell lines.**

ER-negative non-transformed MCF10A and transformed MDA-MB-231 and ER-positive transformed T47D and MCF7 cells were transduced with cKIT^WT^, cKIT^K623M^, or GFP control lentivirus. Transduced cells were FACS sorted and expanded in respective media. MTT viability assays were performed in MCF10A, T47D, MDA-MB-231, and MCF7 cells stably expressing cKIT^WT^, cKIT^K623M^, or GFP controls. Cells were treated with imatinib, dasatinib, dovitinib, or 5-FU for 72 hours. MCF10A-cKIT^WT^ showed response to agents. No impact of cKIT expression on drug sensitivity was observed in established breast cancer models.

**Supplementary Figure S9. Sensitivity of mammary epithelial subpopulations to oxidative stress induced by BSO treatment.**

Growth of sorted Lin^-^EpCAM^low^/CD49f^+^(basal cells) and Lin^-^EpCAM^high^/CD49f^-^ (luminal cells) from patient 1 **(A)** and of Lin^-^MUC1-(basal) and Lin^-^MUC1+ (luminal) epithelial subpopulations sorted from patient 2 **(B)** following 72-hour treatment with increasing concentrations of glutathione depleting agent BSO, measured by MTT assay and normalized to untreated controls. Basal cells showed dose-dependent sensitivity to BSO, while luminal cells were relatively resistant. Note: Both donors are pathogenic BRCA carriers.

