## Supplementary figures and images for "Receptor Tyrosine Kinase Profiling Identifies Chronic Constitutive Floodgate Oxidative Signaling in Glutathione-Independent Human Mammary Luminal Progenitor Cells"

### Supplementary Figures

#### Figure S1

A

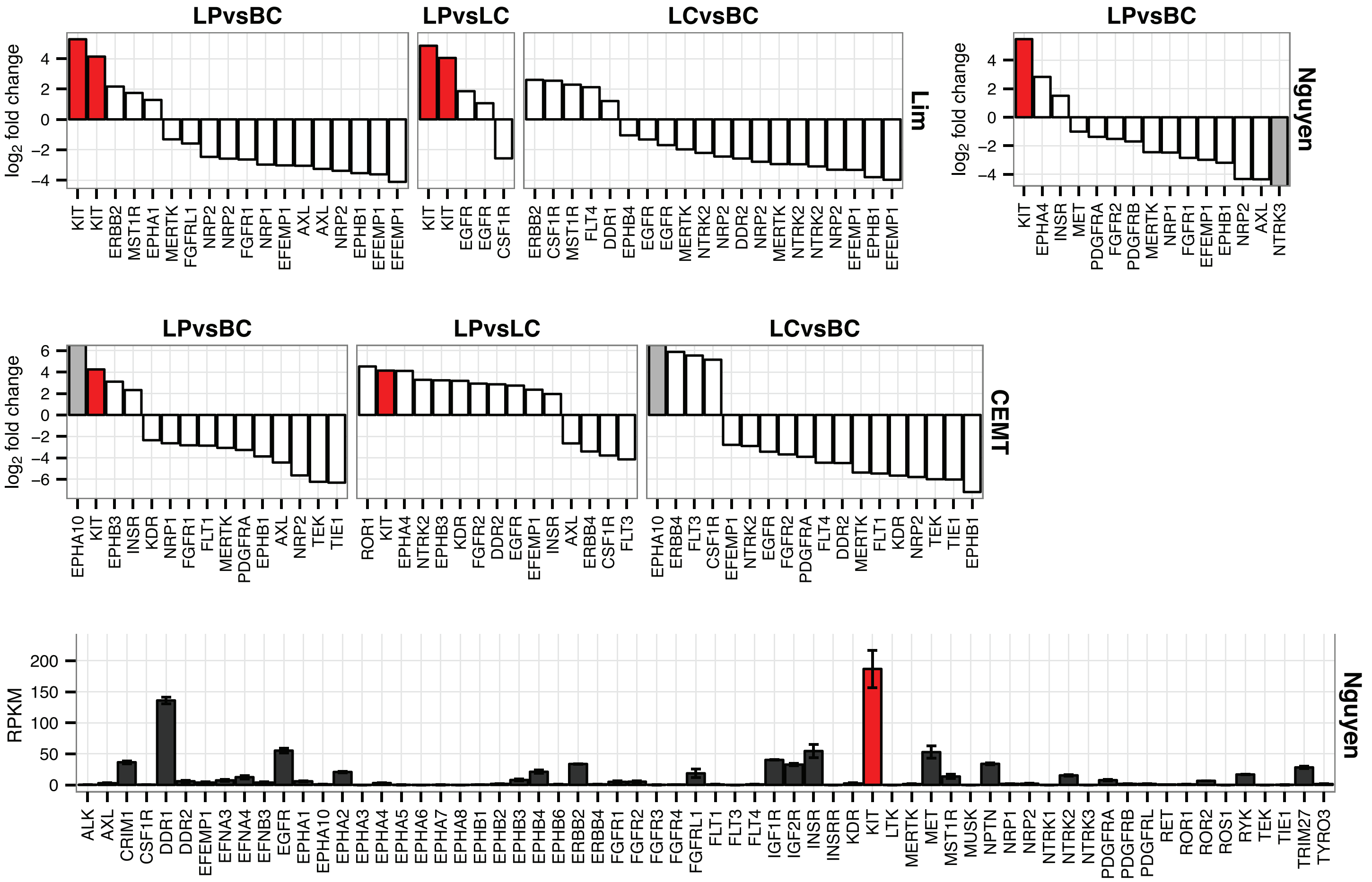


B

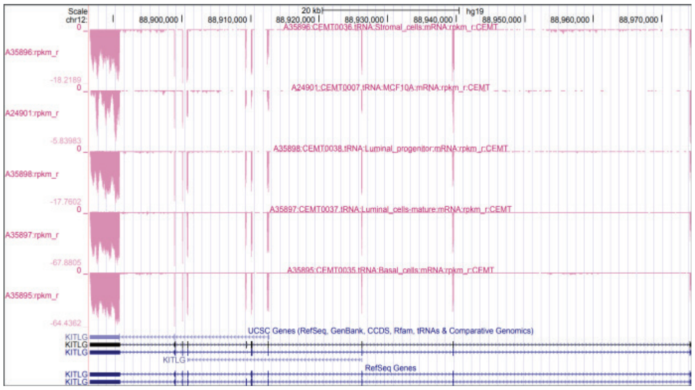

C

Immunohistochemistry - cKIT

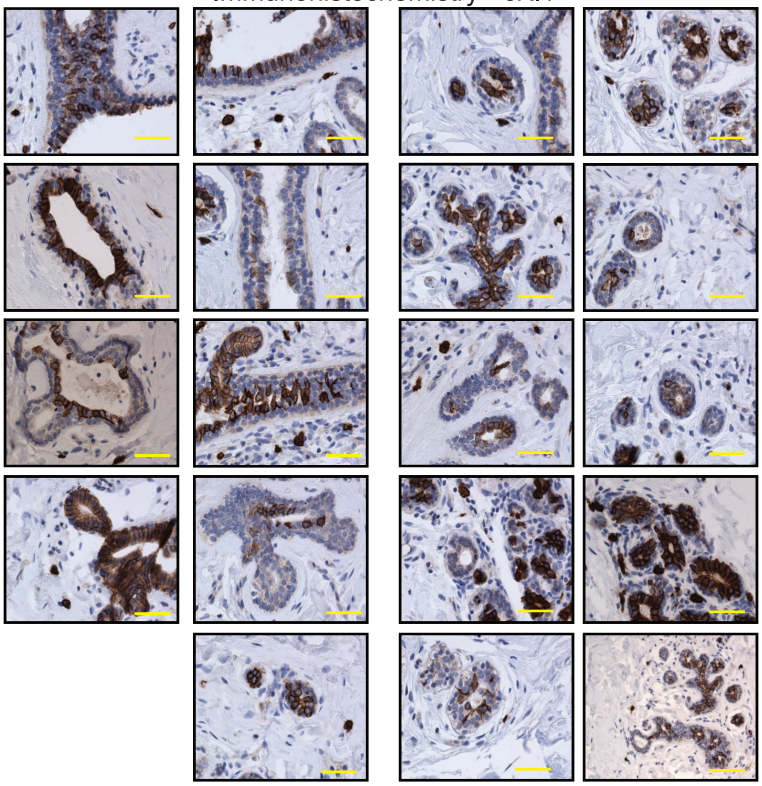

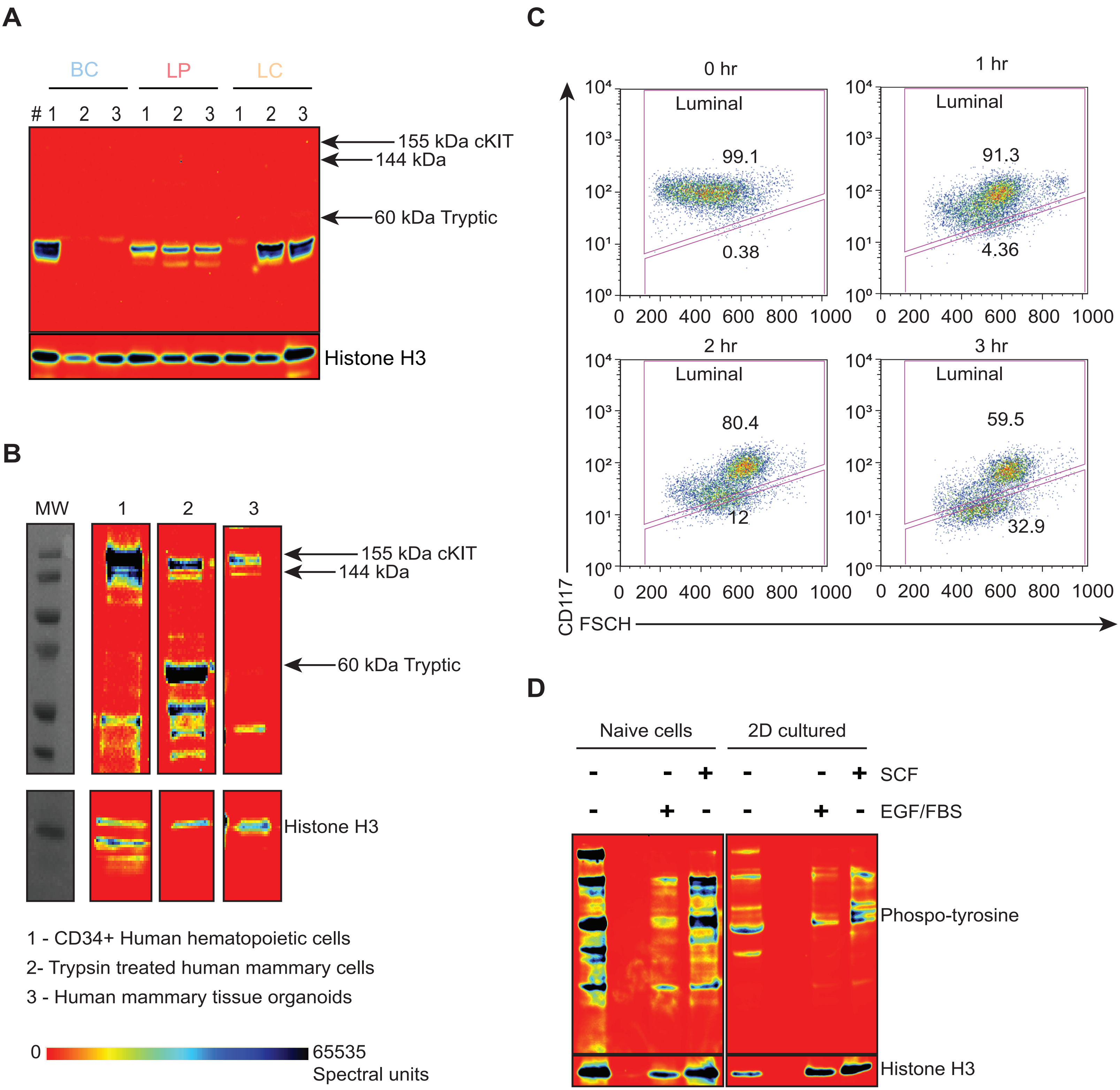

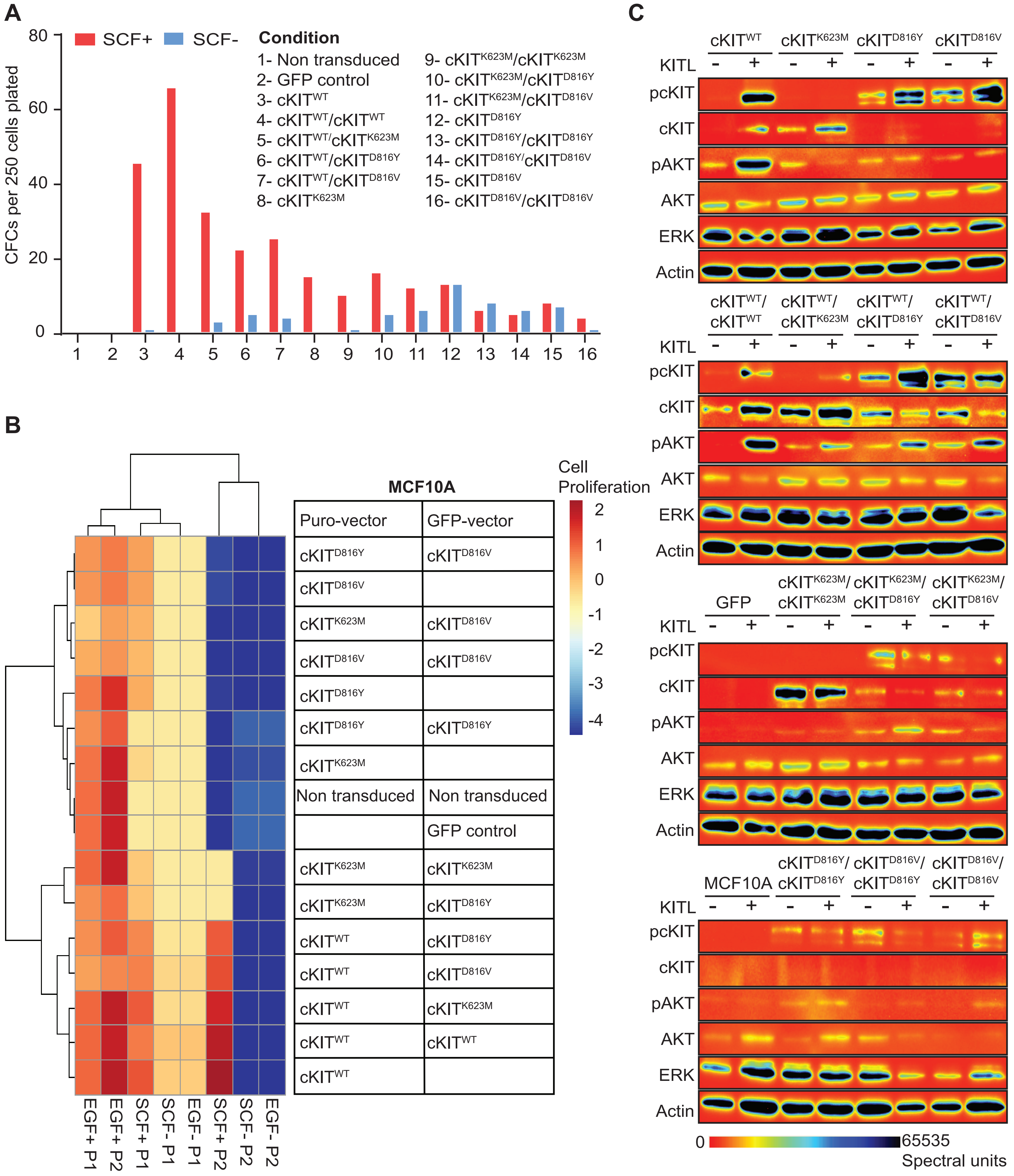

#### Figure S4

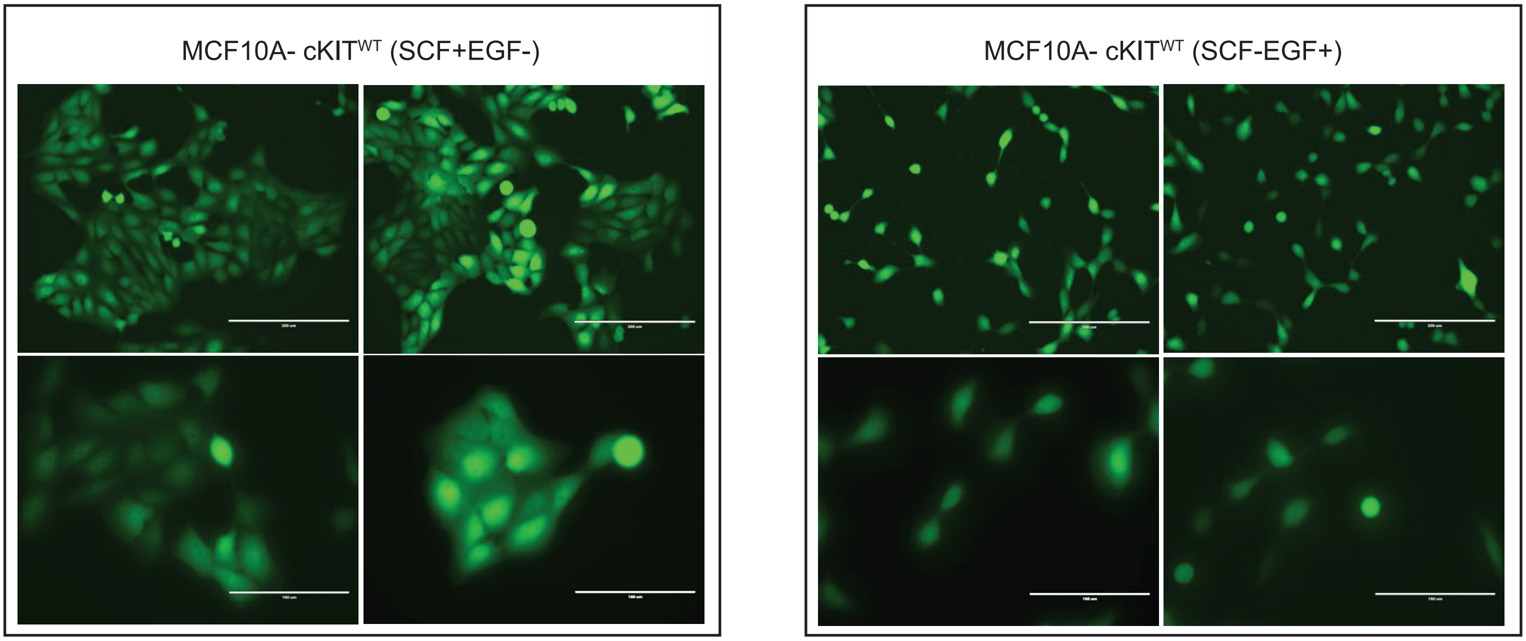

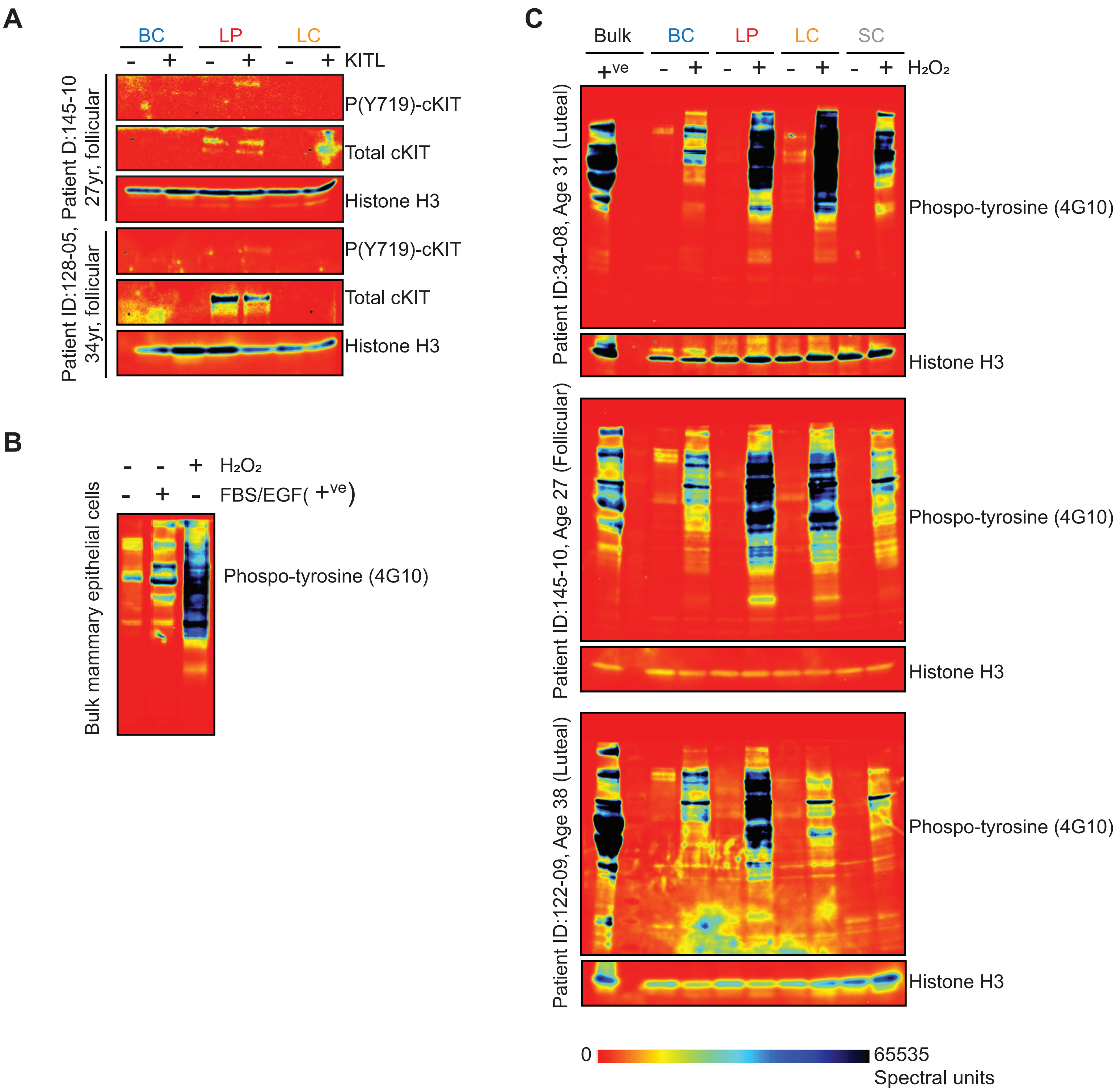

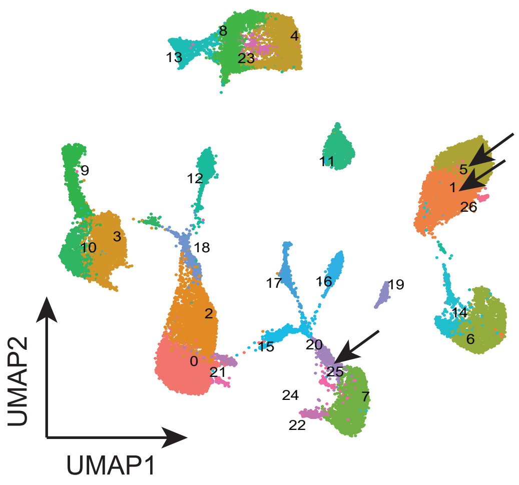

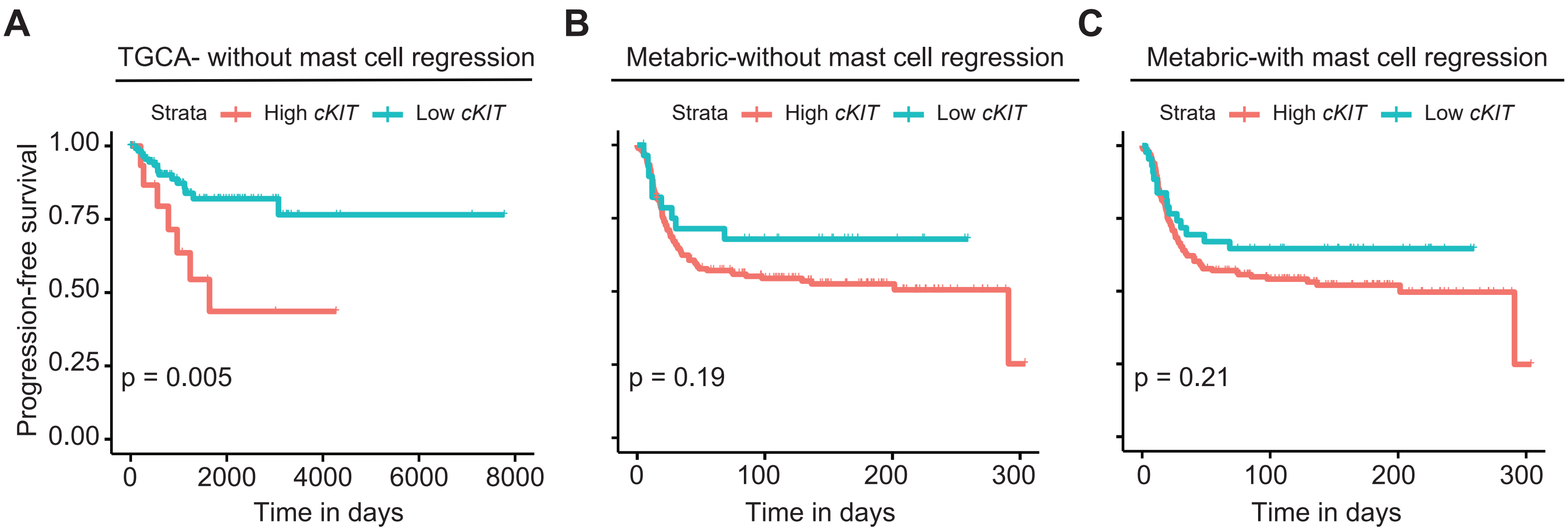

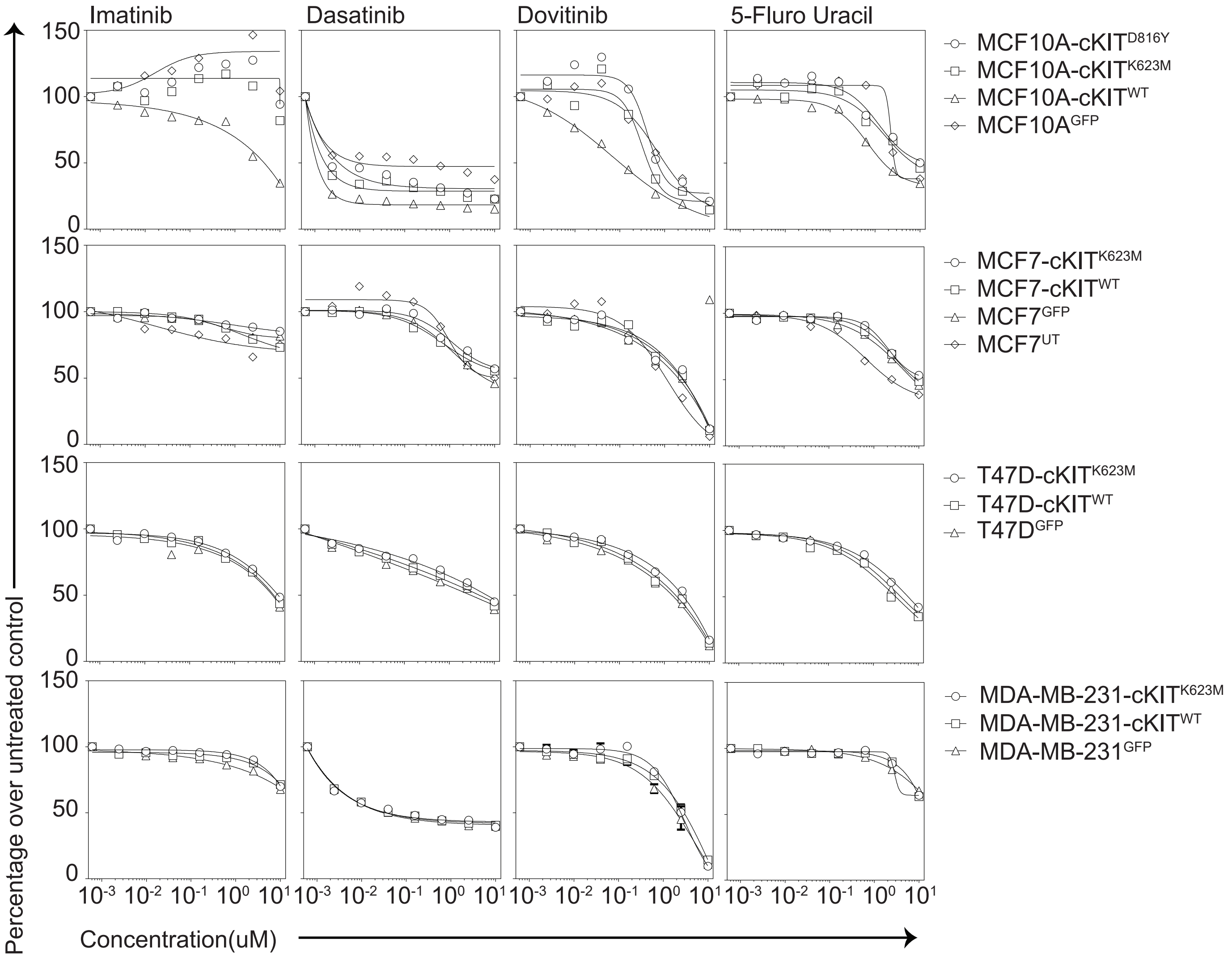

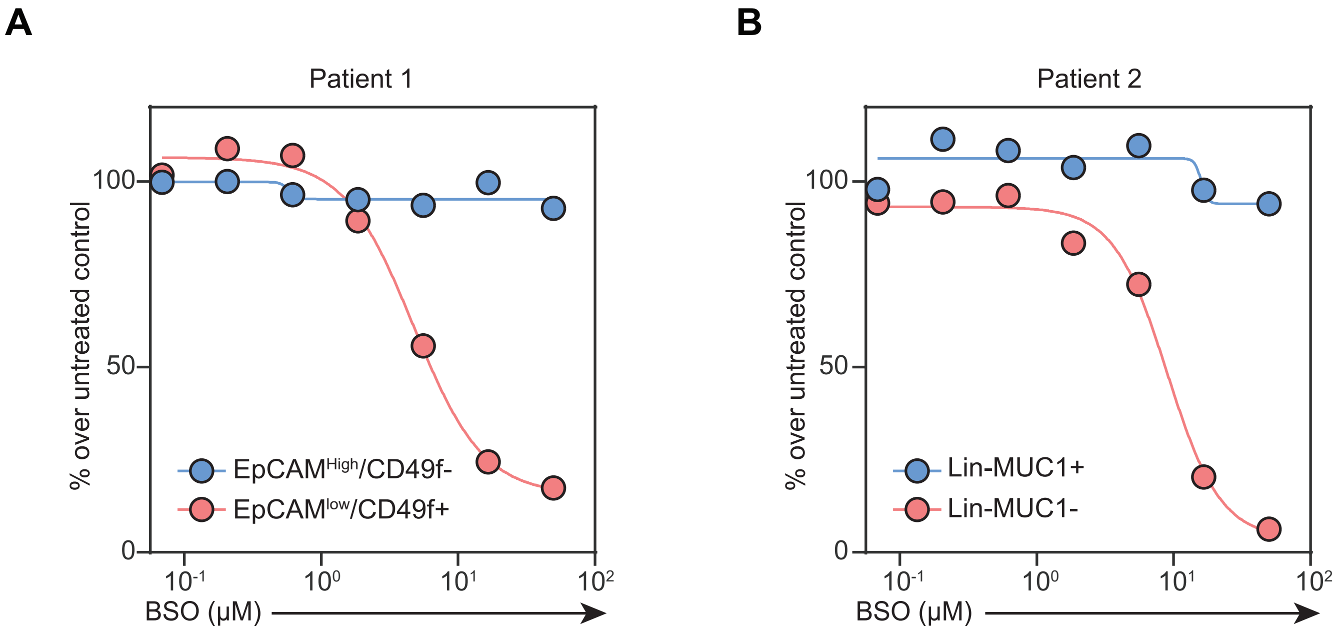
